# Genome-wide temperature-sensitivity of PcG regulation and reduction thereof in temperate *Drosophila melanogaster*

**DOI:** 10.1101/2023.01.02.522482

**Authors:** Susanne Voigt, Christin Froschauer

**Author notes:** SV should be considered as first, senior and corresponding author.

## Abstract

Epigenetic regulation varies with the environment. In the fruit fly *Drosophila melanogaster*, environmental temperature can affect chromatin-based gene regulation. Genes regulated by the Polycomb group (PcG) can vary in their transcriptional output in response to changes in temperature, which typically increases with decreasing temperature. Here, we studied temperature-sensitive expression of PcG target genes on a genome-wide scale, as well as temperature-sensitive enrichment of two histone modifications associated with the regulation of PcG target genes, H3K27me3 and H3K4me3. We investigated temperature-sensitivity in adult flies, and possible differences thereof between populations adapted to temperate and tropical climates. Compared to genes not targeted by the PcG, an elevated number of target genes showed higher expression at the lower temperature, as it is typically observed for PcG regulation. Many of the PcG target genes also exhibited temperature-sensitive H3K4me3 enrichment in the same direction, and the H3K4me3 temperature response correlated positively with that of expression. A small set of target sites also showed temperature-sensitive enrichment of H3K27me3, again with a higher proportion corresponding to increased transcriptional activation at the lower temperature. Overall, higher transcriptional activity at lower temperature was less pronounced in males compared to females, and in temperate compared to tropical flies. Possible trans- and cis-acting factors responsible for reduced expression plasticity in temperate flies were identified, including factors belonging to the Trithorax group (TrxG) and insulator binding proteins, respectively.

## Introduction

The phenotypic outcome of interactions between the genotype and the environment is often mediated by epigenetic changes. Genotypic variation, however, can also affect epigenetic states, as well as the epigenetic response to environmental changes (Cavalli and Heard 2019). In the fruit fly *Drosophila melanogaster*, it is known that the environment, in particular environmental temperature, can have an effect on epigenetic mechanisms (Fauvarque and Dura 1993; Gibert and Peronnet 2021). *D. melanogaster* as a cosmopolitan species has adapted to a wide range of thermal environments (Hoffmann et al. 2003; Ayrinhac et al. 2004; Pool et al. 2017), and is therefore well-suited for studying the interplay between genotypes, epigenetics, and the environment. It has recently spread from its tropical origins in southern-central Africa to nearly all regions around the world including the temperate zones of Europe (David and Capy 1988; Lachaise and Silvain 2004; Stephan and Li 2007; Laurent et al. 2011; Pool et al. 2012; Arguello et al. 2019; Kapopoulou et al. 2020). Temperate climates are characterized by low and varying temperature. Variations in temperature have been shown to affect chromatin-based gene regulation (Fauvarque and Dura 1993). The evolutionary conserved Polycomb group (PcG) of proteins are important epigenetic regulators in *Drosophila* (Kassis and Brown 2013; Simon and Kingston 2013; Steffen and Ringrose 2014; Entrevan et al. 2016; Giner-Laguarda and Vidal 2020; Kuroda et al. 2020) that appear to be sensitive to changes in temperature. Genes regulated by the PcG vary in their transcriptional output in response to changes in the temperature at which flies are reared or held. Expression of PcG-regulated genes typically increases with decreasing temperature (Fauvarque and Dura 1993; Chan et al. 1994; Zink and Paro 1995; Bantignies et al. 2003; Gibert et al. 2011; Voigt et al. 2015). Classically, this has been demonstrated in transgenic assays in which reporter gene expression was controlled by PcG-regulatory sequences derived from prominent PcG target genes. The *miniwhite* reporter for red eye color was mostly used to show temperature-sensitive expression of PcG-regulated genes (Fauvarque and Dura 1993; Chan et al. 1994; Zink and Paro 1995; Bantignies et al. 2003; Gibert et al. 2011). How changes in temperature affect PcG-mediated silencing is still largely unknown, although decreased motion between chromatin domains, in particular PcG-associated domains, which allow long-range chromosomal contacts, have been discussed to contribute to reduced PcG-mediated silencing at lower temperatures (Cheutin and Cavalli 2012; Gibert and Peronnet 2021). In addition, there is also evidence that expression levels of genes encoding for PcG proteins vary with temperature (Voigt et al. 2015; Zhao et al. 2015). PcG proteins modify chromatin by introducing specific histone modifications to epigenetically silence their target genes. They function in multi-protein complexes that are recruited to their target sites by specific cis-regulatory DNA elements called Polycomb response elements (PREs). These act as nucleation sites for PcG complexes to create large repressive Polycomb domains which are characterized by trimethylation of histone 3 at lysine 27 (H3K27me3). PREs consist of different binding sites for proteins that are thought to be involved in PcG recruitment (Kassis and Brown 2013; Simon and Kingston 2013; Steffen and Ringrose 2014; Entrevan et al. 2016; Giner-Laguarda and Vidal 2020; Kuroda et al. 2020). Another group of epigenetic factors, the Trithorax group (TrxG) of proteins, is also involved in the regulation of PcG target genes. PcG and TrxG proteins work in an antagonistic manner to maintain repressed and activated states, respectively (Steffen and Ringrose 2014; Kuroda et al. 2020). More recent evidence suggests a broader function of PcG proteins, in that they not only appear to maintain transcriptional repression, but also to modulate transcription by dampening expression levels of their transcriptionally active target genes (Enderle et al. 2011; Loubiere et al. 2016; Pherson et al. 2017). Many PcG target genes encode for transcription and signaling factors with important roles in development and cell-fate specification, including most prominently the HOX genes (Schuettengruber et al. 2009; Schwartz et al. 2010). In addition to their function in development, PcG proteins appear to be involved in the dynamic regulation of a wide variety of processes such as cell cycle control, spermatogenesis, metabolism, cellular senescence, tissue homeostasis, mitochondrial function, and redox homeostasis (Schuettengruber et al. 2009; Pospisilik et al. 2010; Vidal 2014; Ugrankar et al. 2015). Given their involvement in many essential processes, temperature-induced expression plasticity of PcG target genes might have played a role while adapting to temperate climates. It could have been detrimental by shifting expression levels away from an optimum, and also due to the inability to buffer against temperature fluctuations so that consistent expression levels are maintained across temperatures (Levine and Begun 2008). Indeed, deregulation of PcG genes and the concomitant misexpression of their target genes have been associated with many diseases, including cancer, in humans and also *Drosophila* (Classen et al. 2009; Martinez et al. 2009; Loubiere et al. 2016; Chetverina et al. 2020). Expression plasticity due to temperature, however, could have also been of advantage if changes in expression levels contribute to phenotypic plastic responses that allow the fly to produce the best-suited phenotype in each environmental temperature. Finally, temperature-induced expression plasticity might have also been (nearly) neutral with little effects on fitness (Huang and Agrawal 2016; Mallard et al. 2020).

Here, we studied temperature-sensitive expression of PcG target genes on a genome-wide scale, as well as temperature-sensitive enrichment of the associated repressive histone mark H3K27me3 and the counteracting TrxG-specific histone mark, trimethylation of histone3 at lysine 4 (H3K4me3). We examined temperature-sensitivity in female and male adult *D. melanogaster*, and possible differences thereof between populations adapted to different climates. We found an elevated number of PcG target genes with higher transcriptional activation for expression and enrichment of histone marks at the lower temperature, as it is typically observed for PcG regulation. Overall, higher transcriptional activity at the lower temperature was less pronounced in males compared to females, and in temperate compared to tropical flies.

## Results

### Temperature-sensitive expression of PcG-regulated genes

In order to explore temperature-sensitive expression of PcG-regulated genes, whole-genome expression was assessed in female and male adult flies that were raised at two different temperatures (15° and 28°C). Rearing temperatures were based on the lower and upper ends of the temperature range (14°C to 29°C) in which *D. melanogaster* can successfully complete an entire life cycle (Petavy et al. 2001). We looked at temperature-sensitive expression in flies derived from two populations that are adapted to two contrasting climates: one from temperate France (Lyon) and one from tropical Zambia (Siavonga) (table 1). Due to their prominent function in developmental processes, expression of PcG target genes can be weak in adults. Therefore, in order to capture as much as possible of PcG-regulated expression, we aimed to sequence over 60 million paired-end reads per sample. As a result, the number of uniquely mapped reads ranged from 55,651,302 to 86,514,693 per sample (supplementary table S1), and expression for 758 of the 818 PcG target genes, as defined by Schwartz et al. (2010), could be observed. Principal component analysis (PCA) revealed that most of the variance in PcG target gene expression was explained by sex and rearing temperature (supplementary fig. S1B) with 90% (principal component 1) due to differences between sexes and 4% (principal component 2) due to differences in temperature. Temperature-sensitive expression was observed for the majority of PcG target genes (fig. 1A, supplementary table S4), and was independent of the overall expression levels of the genes (supplementary fig. S3). Compared to nontarget genes (supplementary fig. S4A), PcG targets showed a higher proportion of genes significantly overexpressed at 15°C and a lower proportion of genes significantly overexpressed at 28°C in females (Chi-square tests; proportion of overexpression at 15°C: France *P* < 0.0001 & Zambia *P* < 0.0001; proportion of overexpression at 28°C: France *P* < 0.0001 & Zambia *P* < 0.0001) and males, though this was not significant in French males (Chi-square tests; proportion of overexpression at 15°C: France *P* < 0.05 & Zambia *P* < 0.0001; proportion of overexpression at 28°C: France *P* = 0.25 & Zambia *P* < 0.01). The observed directionality of temperature-sensitive expression of PcG target genes, therefore, fits the typical temperature sensitivity of PcG regulation with higher expression at lower temperatures (Fauvarque and Dura 1993; Chan et al. 1994; Zink and Paro 1995; Bantignies et al. 2003; Gibert et al. 2011; Voigt et al. 2015; Voigt et al. 2019). As it was observed before (Voigt and Kost 2021), this was more pronounced in females than in males. In response to unfavorable conditions, females sometimes, however, retain their mated eggs (Horváth and Kalinka 2018). Since many PcG targets function in development, and developmental time also varies with temperature (Trotta et al. 2006), the observed patterns of temperature-dependent expression could also arise due to females retaining their eggs. Additionally, ovary size is known to vary with temperature (Delpuech et al. 1995). Thus, in order to rule out these two factors as the main reasons for the observed PcG-typical temperature response, we looked separately at the expression of PcG targets with high expression in ovaries and embryos, and at those with no expression in both (Gramates et al. 2022). As a result, PcG targets with no expression in ovaries and embryos showed a much larger proportion of overexpression at 15°C, while for those with high expression in both, the proportion of overexpression at 28°C was much larger (supplementary fig. S4B) (Chi-square tests; proportion of overexpression at 15°C: France *P*<0.01 & Zambia *P*<0.001; proportion of overexpression at 28°C: France *P*=0.11 & Zambia *P*<0.001). Furthermore, the large majority of PcG targets overexpressed at 15°C in males, also exhibited the same overexpression in females (supplementary fig. S5). Overrepresentation analyses (ORA) of gene ontology (GO) terms were then performed to uncover biological processes in which PcG targets with temperature-sensitive expression are involved. Analyses were run against a background sample comprising all expressed PcG targets to which the sets of PcG targets with overexpression at the specific temperature in at least one of the populations were compared. Significantly overrepresented GO terms (FDR < 20%) were only observed for females (supplementary tables S10-S15) with the majority of overrepresented terms for PcG targets overexpressed at 15°C involved in developmental processes (fig. 1E). While for those PcG targets overexpressed at 28°C overrepresented GO terms mostly included such involved in reproductive processes which correlates well with a general increase in reproduction at warmer temperatures (Trotta et al. 2006).

**Fig. 1.**
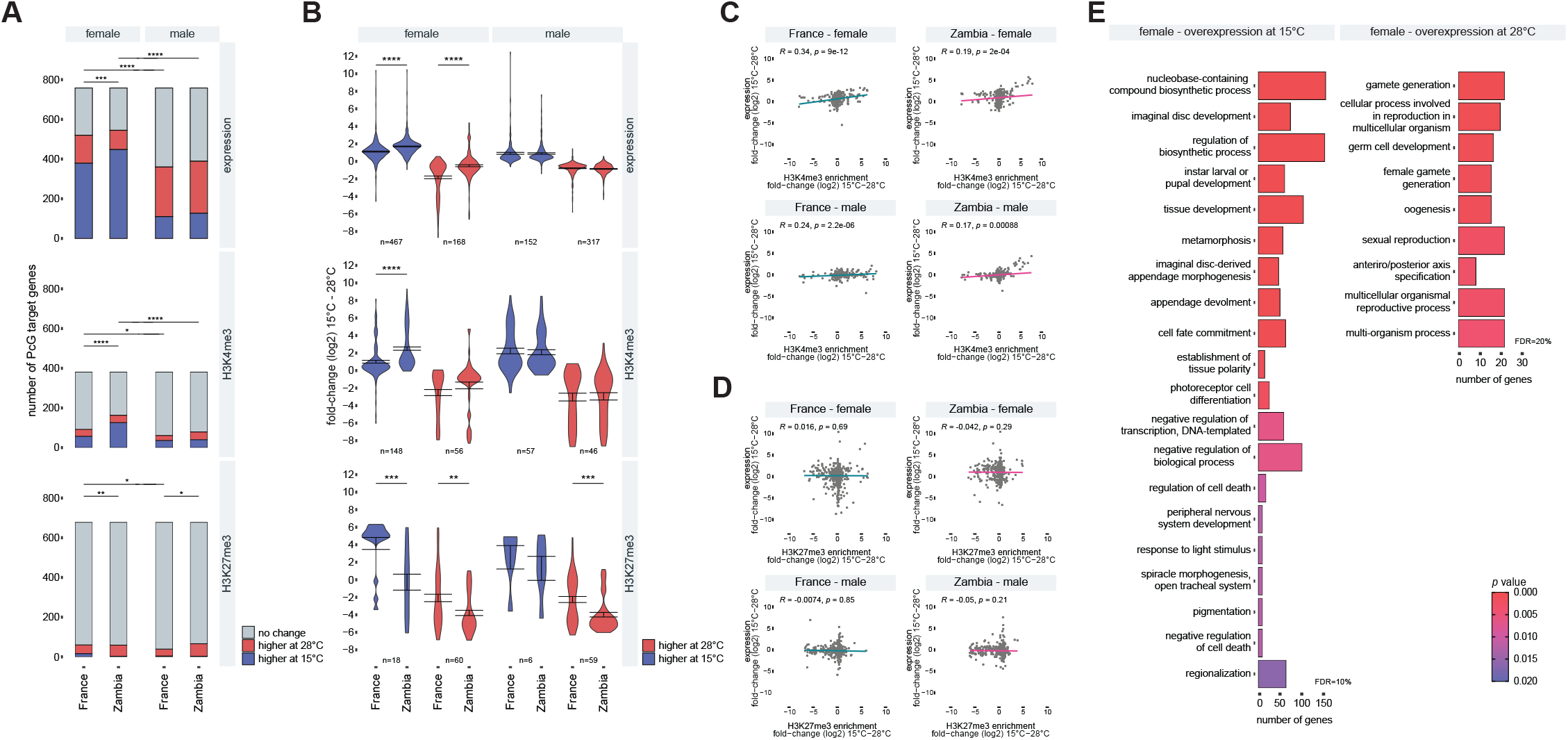
Temperature-sensitivity of PcG-regulated genes. (A) Proportions of PcG target genes with overexpression at 15°C (blue) and 28°C (red) rearing temperature in adult flies of the temperate population sample from France and the tropical sample from the ancestral species-range in Zambia, as well as of those targets associated with overenrichment at 15°C (blue) and 28° (red) of TrxG/PcG-associated histone modifications H3K4me3 and H3K27me3. Differences were assessed using Chi-square tests. (B) Mean 15°/28°C foldchanges of PcG targets overexpressed at 15°C (blue) and 28°C (red), as well as H3K4me3 and H3K27me3 peaks at PcG targets with overenrichment at 15°C (blue) and 28°C (red). n corresponds to the number of PcG targets or peaks associated with PcG targets exhibiting temperature-sensitive expression or enrichment. Differences between population samples were assessed with t-tests. (C) Correlation between temperature responses (i.e., 15°/28°C foldchanges) of PcG target gene expression and associated histone modifications. Spearman’s rank correlation coefficient (R) was estimated. (E) Gene ontology overrepresentation analysis for biological processes in which PcG targets with temperature-sensitive expression are involved. Analyses with significant results at an FDR of at least 20% are shown. * *P* < 0.05; ** *P* < 0.01; *** *P* < 0.001; **** *P* < 0.0001.

**Table 1.**
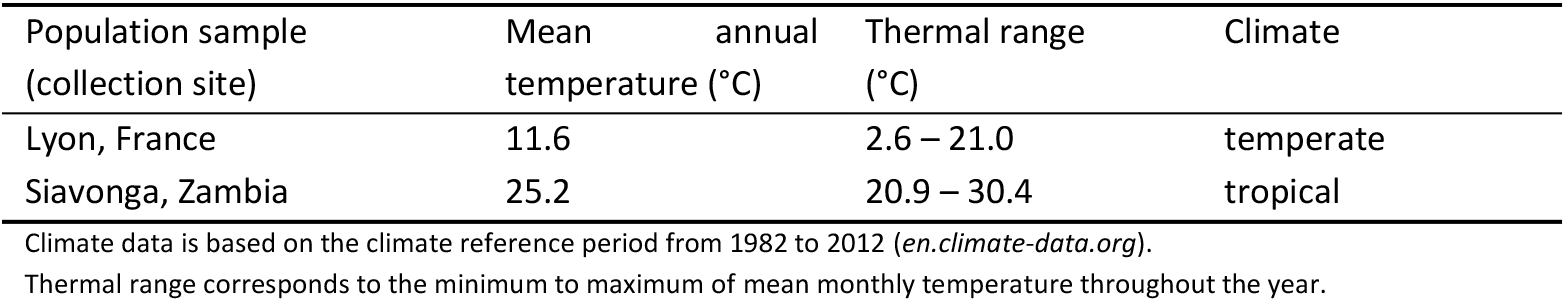
Origins of population samples and climate of origins.

### Differences in temperature-sensitive expression of PcG-regulated genes between temperate and tropical flies

For nontarget genes, the proportion of temperature-sensitively expressed genes was increased in Zambian compared to French flies for both females and males (supplementary fig. S4A). This increased temperature plasticity of expression in tropical flies is consistent with the results of earlier studies that compared temperate and tropical fly populations from Australia (Levine et al. 2011) and North America (Zhao et al. 2015). Nontarget genes exhibited a similar proportion of genes overexpressed at 15 °C in the two population samples, whereas the proportion of genes overexpressed at 28°C was higher in tropical Zambian flies for both sexes (Chi-square tests; proportion of overexpression at 15°C: female *P* = 0.88 & male *P* = 0.52; proportion of overexpression at 28°C: female *P* < 0.0001 & male *P* < 0.0001). In contrast, overall plasticity for PcG target gene expression was not significantly different between French and Zambian flies. In females, both the proportions of genes overexpressed at 15°C and those of genes overexpressed at 28°C, however, significantly differed for PcG targets between the two population samples. While the proportion of overexpression at 28°C was increased in the temperate samples, a higher proportion of overexpression at 15°C was observed for the tropical sample (Chi-square tests; proportion of overexpression at 15°C: female *P* < 0.001 & male *P* = 0.26; proportion of overexpression at 28°C: female *P* < 0.01 & male *P* = 0.55). This is also seen for the overall foldchanges between temperatures of PcG target genes with overexpression at one temperature. In females, Zambian PcG target gene expression showed a higher overexpression at 15°C, while French expression had a higher overexpression at 28°C (fig. 1B). Interaction effects between population and rearing temperature were assessed for each gene to further explore the differences in the temperature response of PcG target gene expression between temperate and tropical flies. Again, this difference was more prevalent in females than in males. At an FDR of 5%, 22 male PcG targets showed significant population x temperature interactions, while in females 208 PcG targets exhibited such interaction effects (table S7). A large majority of the PcG targets with significant interaction effects in females showed a higher overexpression at 15°C in Zambian compared to French flies (fig. 2A). Although to a lesser extent, there were also significantly more PcG targets with significant population × temperature interactions more overexpressed at 28°C in French compared to Zambian flies (fig. 2A). Taken together, females not only showed higher overexpression of PcG targets at 15°C in general, but also more differences in the temperature response of PcG target gene expression between temperate and tropical flies than males. This was most obvious in the reduction of expression plasticity of PcG targets overexpressed at 15 °C, with temperate female flies showing less overexpression at the lower temperature. In most of these cases, expression was weaker at 15°C in French than in Zambian flies, though increased expression at 28°C also contributed to the reduced 15°C/28°C expression ratios observed in temperate flies (supplementary fig. S6A). The sets of PcG target genes with overexpression at one temperature in one population sample, as well as those with equal overexpression at one temperature in both populations for both sexes and their overlaps were assessed for overrepresented GO terms (biological processes). Overrepresentation was again estimated against the background sample of all expressed PcG targets (supplementary tables S16-S33). Many of the GO terms overrepresented for PcG targets more highly overexpressed at 15°C in Zambian females were involved in developmental processes, while most GO terms overrepresented for targets more highly overexpressed at 28°C in Zambian females were associated with reproductive functions (fig. 2B). The latter is consistent with the observation of tropical flies having a higher reproductive productivity at warmer temperatures than temperate flies (Trotta et al. 2006). Although to a much lesser extent, greater overexpression at 15°C was also observed for French flies with five PcG target genes shared between the sexes (fig. 2A and supplementary fig. S6C). Two of these genes encode for enzymes active in mitochondria (*SlgA* and *Prx2540-2*) (Gramates et al. 2022), and two encode for myokines (*Obp99b* and *Amyrel*) that are upregulated due stress (Rai et al. 2021). *Prx2540*-2 encodes for a peroxidase with functions in oxidative stress response, and as well as the product of *Amyrel*, it was shown to have neuroprotective effects and to be involved in healthy aging (Detienne et al. 2018; Rai et al. 2021). This might be related to that, compared to tropical flies, temperate flies are thought to be better at dealing with oxidative stress (González et al. 2009; Guio et al. 2014; Ramnarine et al. 2022).

**Fig. 2.**
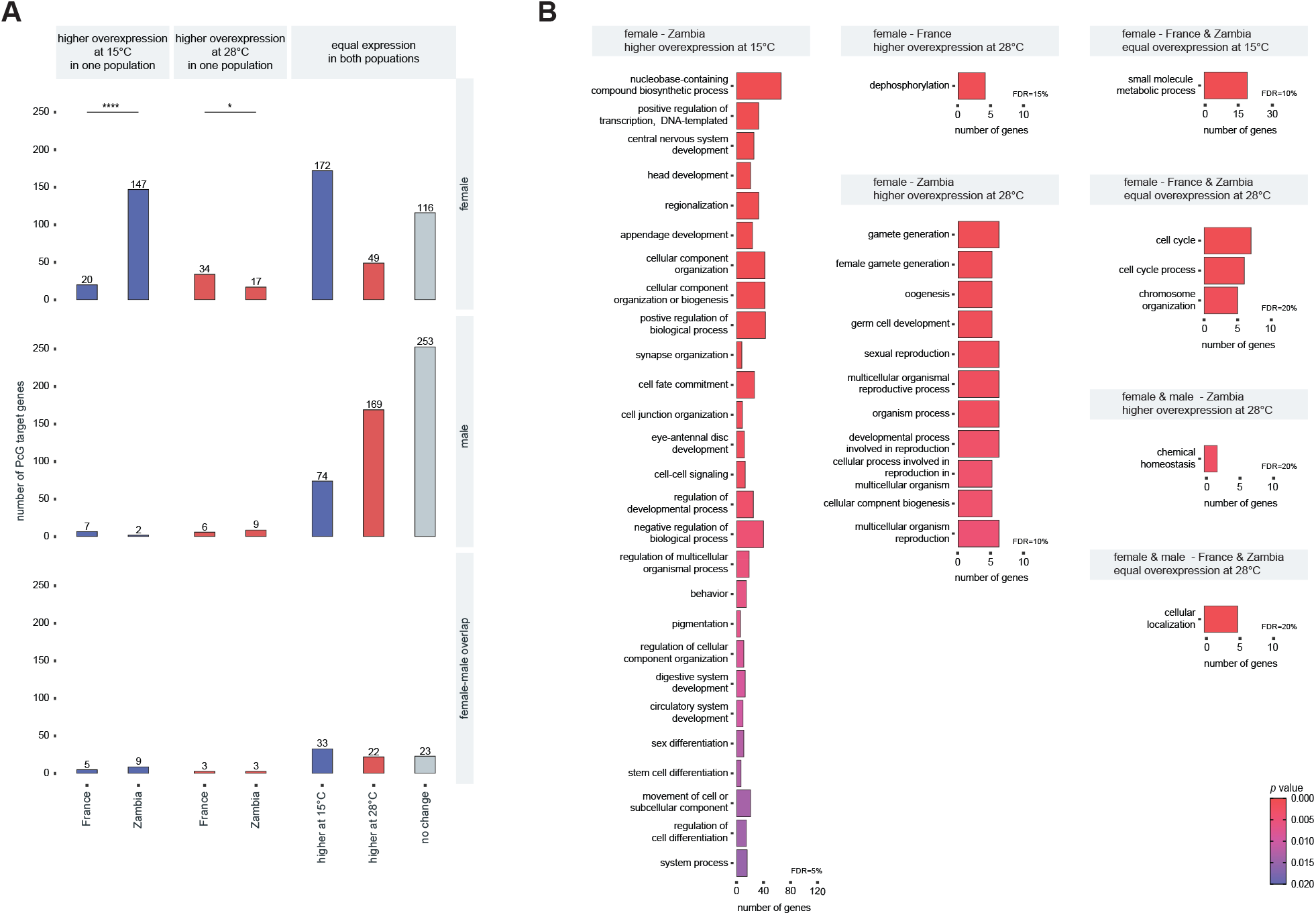
Differences in temperature-sensitive PcG target gene expression between temperate French and tropical Zambian flies. (A) Number of PcG target genes with significant population x temperature interaction effects in their expression (FDR < 5% and with a less conservative cutoff of P < 0.05 for female-male overlaps) and higher overexpression at 15°C or 28°C in one of the population samples, as well as of those with similar expression patterns across temperatures in temperate and tropical flies. Differences in number were assessed using Chi-square tests. (B) Gene ontology overrepresentation analysis for biological processes in which PcG targets with different or equal temperature-sensitive expression patterns in temperate and tropical flies are involved. Analyses with significant results at an FDR of at least 20% are shown. * *P* < 0.05; **** *P* < 0.0001.

### Temperature-sensitive enrichment of histone modifications at PcG target genes

As for expression, temperature-sensitive enrichment of histone modifications in PcG target gene regions was assessed in genome-wide ChIP-seq assays in adult flies of both sexes that were reared at 15°C and 28°C. For the two population samples, the repressive histone mark typical for PcG regulation H3K27me3 (Kassis and Brown 2013; Simon and Kingston 2013; Steffen and Ringrose 2014; Entrevan et al. 2016; Giner-Laguarda and Vidal 2020; Kuroda et al. 2020), as well as H3K4me3 typical for transcriptional activation conferred by the TrxG (Schuettengruber et al. 2011; Piunti and Shilatifard 2016; Chetverina et al. 2020) were assayed. Over 25 million paired-end reads per sample were sequenced (supplementary table S2). Pearson’s correlation of genome-wide binding intensity among biological replicates was high, and ranged between 0.949 and 0.997 (supplementary table S3). Global enrichment profiles were characteristic for each histone modification and similar among samples (supplementary figs. S2B and S2C). We obtained 780 H3K27me3 and 483 H3K4me3 peaks associated with PcG target genes, which corresponds to 678 and 381 of the 818 PcG target genes identified by Schwartz et al. (2010), respectively (supplementary tables S4 and S5). The majority of PcG targets, therefore, showed PcG-typical H3K27me3 enrichment. Due to the averaging effect across all body parts, many PcG targets were associated with enrichment of both H3K4me3 and H3K27me3 (supplementary fig. S1A). Nonetheless, expression levels of each sample were positively correlated with H3K4me3 enrichment levels and negatively correlated with H3K27me3 levels (supplementary figs. S2A and S7), which is consistent with the characteristic effect of each histone modification on transcription. PCA results for both histone modifications were similar to those observed for expression with most of the variance explained by sex differences and differences in rearing temperature (supplementary figs S1C and S1D). As for expression, temperature-sensitivity of H3K4me3 enrichment at PcG targets differed between the sexes with females having more PcG targets associated with changes in enrichment due to temperature (fig. 1A). Furthermore, more PcG targets harbored increased H3K4me3 enrichment at 15°C than at 28°C in females (Chi-square tests; female: France *P* < 0.05 & Zambia *P* < 0.0001; male: *P* = 0.18 & Zambia *P* = 1), which corresponds to the observed increased transcriptional activation of PcG targets at 15°C. The temperature response of H3K4me3 enrichment was significantly correlated to that of expression in both sexes (fig. 1C). H3K4me3 enrichment at PcG targets, therefore, tends to change in the same direction due to temperature as expression. Although expression and enrichment levels were significantly correlated (supplementary fig. S7), overall, no significant correlation in the temperature response could be observed between expression and H3K27me3 enrichment (fig. 1C). This might be primarily attributable to the lower proportion of temperature-sensitive H3K27me3 enrichment, but also the averaging effect across whole flies and the nature of H3K27me3 as a repressive mark might play a role. Temperature-sensitivity of H3K27me3 for both sexes, however, consisted in large parts of higher enrichment at 28°C than at 15°C (Chi-square tests; female: France *P* < 0.001 & Zambia *P* < 0.0001; male: *P* < 0.0001 & Zambia *P* < 0.0001). As H3K27me3 is associated with transcriptional repression, this is consistent with increased transcriptional activity at lower temperatures as typical for PcG regulation. Furthermore, French flies exhibited fewer PcG targets associated with higher H3K27me3 enrichment at 28°C relative to 15°C than Zambian flies, with a significantly lower number in males (fig. 1A) (Chi-square tests; proportion of overexpression at 28°C: female *P* = 0.25 & male *P* < 0.01). Similarly, overall 15°C/28°C foldchanges of peaks with increased H3K27me3 enrichment at 28°C were higher in Zambian flies for both sexes (fig. 1B). In females, a higher number of PcG targets was associated with increased H3K27me3 enrichment at 15°C (Chi-square tests; proportion of overexpression at 15°C: female *P* < 0.01 & male *P* = 1), as well as a higher mean 15°C/28°C foldchange for peaks with increased H3K27me3 enrichment at 15°C in French compared to Zambian flies (figs. 1A and 1B). This might indicate less repression of PcG targets at 28°C compared to 15°C, which matches the higher proportion of PcG targets with increased expression at 28°C in French female flies (fig. 1A). While no significant differences in increased H3K4me3 enrichment at 28°C relative to 15°C were observed between the two population samples, a higher number of PcG targets associated with increased H3K4me3 enrichment at 15°C was observed in Zambian compared to French females (Chi-square tests; proportion of overexpression at 15°C: female *P* < 0.0001 & male *P* = 0.71, proportion of overexpression at 28°C: female *P* = 0.9 & male *P* = 0.09) with a corresponding increased mean foldchange for peaks with increased H3K4me3 enrichment at 15°C in Zambian females (figs. 1A and 1B). This indicates higher transcriptional activity of PcG targets at 15°C in Zambian flies, similar to what was observed for PcG target gene expression.

### Candidates for trans- and cis-acting factors responsible for differences in temperature-sensitive PcG target gene expression between temperate and tropical flies

Both trans-regulatory and cis-regulatory changes might be responsible for the different temperature responses of PcG targets that were observed between temperate French and tropical Zambian flies. While differences in cis-acting factors would directly affect target gene expression through changes in regulatory elements, differences in trans-acting factors would involve activity changes of factors that regulate target gene expression. Expression of PcG target genes is regulated by PcG and TrxG proteins, which makes them likely candidates for trans-acting factors responsible for changes in the transcriptional response to temperature. In addition, insulator proteins, which are involved in chromatin demarcating and genomic architecture, were also shown to be of importance in regulating PcG target gene expression and ensuring PcG-mediated repression (Sigrist and Pirrotta 1997; Comet et al. 2011; Li et al. 2011; Li et al. 2013; Erokhin et al. 2021). We thus scanned these trans-acting factors (supplementary table S34) for differences in their transcriptional response to temperature between French and Zambian flies. While no significant interaction effects were found in males, expression of genes encoding for some of these factors exhibited significant interactions between population and rearing temperature (FDR < 5%) in females (fig. 3A). These included one gene encoding for the PcG recruiter protein Grh, one encoding for the insulator protein Elba3, and seven genes encoding for products classified as TrxG proteins (fig. 3A). Most PcG targets that were observed having population x temperature interactions in their expression, showed less differences between temperatures in the French population sample, often with lower expression at 15°C in French compared to Zambian flies (supplementary fig. S6A). Consistent to that, nearly all of the considered trans-acting factors with significant interaction effects harbored less expression differences between temperatures in French flies. However, counterintuitively, expression of the PcG recruiter gene *grh* was higher at 15°C in Zambian flies compared to French flies, whereas the genes encoding for TrxG proteins were more expressed at 15°C in French flies. The exact mechanism of PcG recruitment to their target genes, however, is still poorly understood. Although Grh is known to be involved in PcG recruitment (Blastyák et al. 2006), similar to other classified PcG recruiter proteins, it also appears to participate in transcriptional activation and TrxG function (Bray and Kafatos 1991; Hur et al. 2002). Two of the genes with significant interaction effects and classified to encode for TrxG proteins (Gramates et al. 2022), *simj* and *Mi-2*, actually encode for subunits of the chromatin-remodeling NURD complex, whose activity is thought rather to facilitate PcG repression (Bornelöv et al. 2018). Other confounding factors arise from the averaging effect across the whole organism, as well as that there was a wider variety of changes in temperature responses of PcG targets with significant population x temperature interaction effects. Still given the significant interactions in their expression between population and temperature, and the lower differences in expression levels between temperatures in French flies, those trans-acting factors are likely candidates involved in mediating reduced temperature-induced plasticity of PcG target gene expression in temperate flies.

**Fig. 3.**
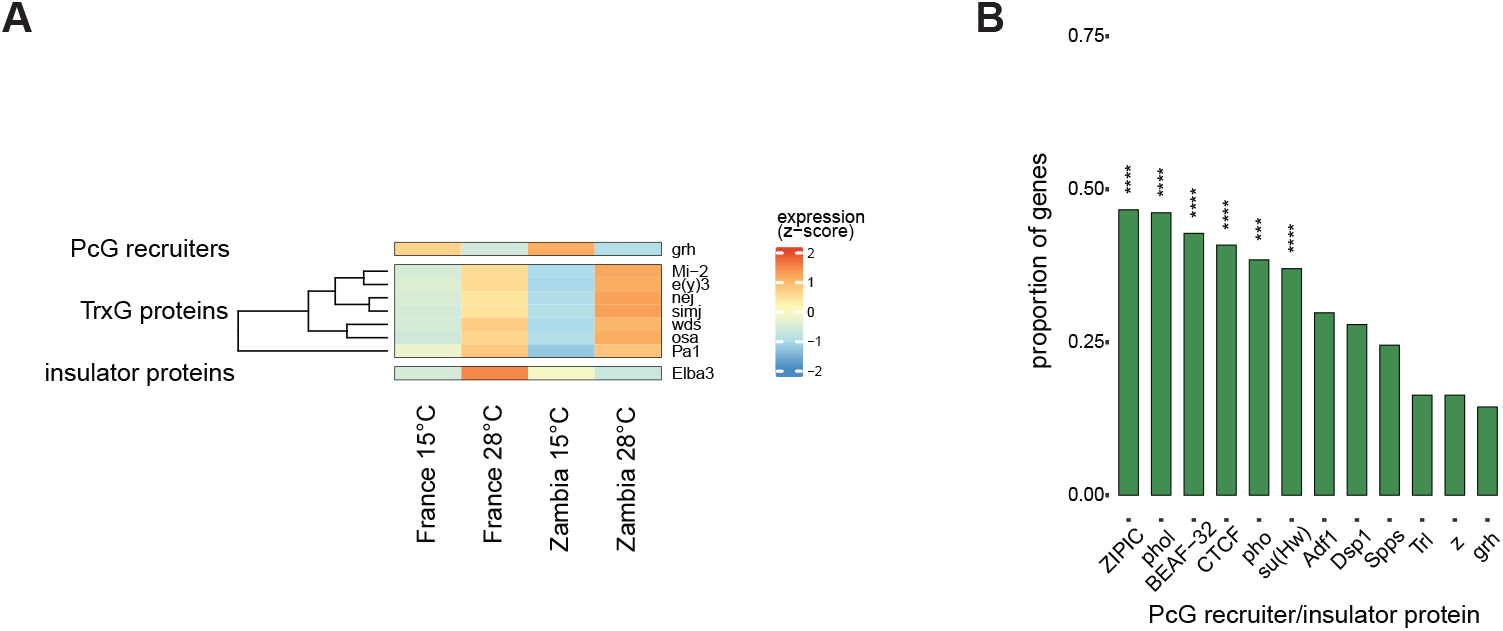
Candidates for cis- and trans-acting factors involved in differences in temperature-sensitive PcG target gene expression between temperate and tropical female flies. (A) Expression levels across temperatures of genes with PcG/TrxG-related functions and significant population x temperature interaction effects in their expression (FDR < 5%). Gene-wise standardized expression levels (Z scores) are given. (B) Proportions of PcG target genes with significant population x temperature interactions in their expression that harbor significant differentiation of DNA binding motifs of PcG recruiter and insulator proteins between the populations in their gene regions. Asterisks correspond to significant overrepresentation (FDR < 5%) of differentiated motifs at PcG target genes with significant interaction effects compared to those with none. *** *P* < 0.001; **** *P* < 0.0001.

Cis-regulatory elements might also harbor differences leading to changes in temperature-sensitivity of PcG target gene expression between temperate and tropical flies. We, therefore, searched for highly differentiated SNPs between the populations that led to motif changes of transcription factor binding sites in the gene regions of PcG targets with significant population x temperature interactions in their expression. For this, we calculated genome-wide *F_ST_* per SNP (Weir and Cockerham 1984) using sequence data from the Drosophila Genome Nexus (DGN) resource for 96 and 197 genomes from the French and the Zambian populations. This data set includes genomes of the same populations as used in this study, as well as the genomes of the studied fly lines. For each chromosome arms, SNPs with *F_ST_* values above the 99^th^ quantile were considered as significantly differentiated between French and Zambian flies (see supplementary file 1). Motif differentiation between the two populations was then calculated for fragments of ±20bp around the outlier SNPs for 591 available binding motifs corresponding to binding sites of 313 different transcription factors (supplementary table S35). Cutoff values for significant motif differentiation were based on the 97.5^th^ quantiles of the empirical distributions of differentiation scores for each motif that were estimated from 10,000 randomly drawn *F_ST_* outlier SNPs located within ±2kb gene regions along the genome. Motifs with low differentiation were excluded from the analysis (see supplementary file 1). Significantly differentiated motifs that were also overrepresented in gene regions of PcG targets with significant population x temperature interactions in their expression were considered as candidates for cis-acting changes. Over-representation was assessed against a background sample of significant scores in gene regions of PcG targets lacking significant interaction effects. This yielded, after filtering for factors with observed expression in each sex, 118 and 18 transcription factors associated with changes in temperature-sensitive PcG target gene expression between French and Zambian flies for females and males, respectively, of which 17 were shared between the sexes (supplementary figs. S8A-C, supplementary tables S36-37). Of the DNA-binding proteins associated with PcG regulation (i.e, PcG recruiter and insulator proteins), significant changes in motifs of insulator proteins were predominately observed as over-represented (fig. 3B and supplementary fig. S8B). While for males only CTCF showed overrepresented motif changes in PcG target gene regions with population x temperature interaction effects, nearly all insulator proteins with available motifs (supplementary table S35) were overrepresented in females. These included changes in motifs of CTCF, Su(Hw), BEAF-32 and ZIPIC, but not motif changes of the GAGA factor, which is encoded by *Trl* and associated with both insulator and PcG recruiter function (fig. 3B). For females, significant motif changes between temperate and tropical populations were also overrepresented for the PcG proteins Pho and Phol (fig. 3B), which have important roles in PcG recruitment (Wang et al. 2004; Wang et al. 2010; Brown et al. 2018).

## Discussion

The characteristic temperature-sensitivity of PcG-regulated genes with increased transcription at lower temperatures was discovered early on, and demonstrated several times by reporter gene assays using PRE sequences from prominent PcG targets (Fauvarque and Dura 1993; Chan et al. 1994; Zink and Paro 1995; Bantignies et al. 2003). Here, we investigated temperature-sensitivity of PcG regulation on a genome-wide scale in the natural genomic context of the genes. For this, we examined expression levels of all classified PcG target genes, as well as enrichment levels of associated histone modifications in adult flies reared at a low and a high temperature. In order to assess how temperature-sensitivity might have changed while adapting to temperate environments, flies were derived from two different populations, one temperate from France and one tropical from the ancestral species range in Zambia. An elevated number of PcG target genes compared to nontarget genes showed indeed PcG-typical overexpression at the lower temperature. As observed before (Voigt and Kost 2021), this overexpression at 15°C was more pronounced in females than in males. One explanation for this difference might be temperature-induced changes in reproductive output as well as in ovary size (Delpuech et al. 1995; Trotta et al. 2006), and a relatively larger size of female reproductive organs within the body. However, PcG targets overexpressed at 28°C were predominantly involved in reproductive processes, while those overexpressed at 15°C were more involved in developmental processes. Another possible explanation might be changes in developmental time in response to temperature (Trotta et al. 2006), if females would retain their eggs due to less suitable temperatures. Yet such blocking of egg laying is mostly observed when oviposition and nutritive sites are limited (Boulétreau-Merle and Terrier 1986), which was not the case in this experiment. Furthermore, the proportion of PcG targets with higher transcription at 15°C was higher for those with no expression in ovaries and embryos compared to those with high expression in both, for which an elevated proportion of targets with overexpression at 28°C was observed instead. Since this study, investigated temperature-induced changes in whole flies, changes in expression levels in response to temperature might also arise from differences in the scaling relationships between the different organs of the body. The sizes of organs can change with rearing temperature, and that change in size can be stronger for one organ than the other (Shingleton et al. 2009). This alone could therefore result in overall expression changes between flies reared at different temperatures. An earlier study, however, also observed PcG-typical overexpression at lower temperature in specific tissues for several PcG target genes using the same population samples (Voigt and Kost 2021). Furthermore, classic experiments have often demonstrated temperature-sensitivity typical for PcG regulation using red-eye color as a marker (Fauvarque and Dura 1993; Chan et al. 1994; Zink and Paro 1995; Bantignies et al. 2003). Thus, regulatory changes are most likely involved in causing increased transcription at lower temperature of PcG-regulated genes.

Overexpression at lower temperature was also significantly more pronounced in the tropical population sample from Zambia than in the temperate from France. French female flies exhibited less overexpression at 15°C, but also, though to a lesser extent, more overexpression at 28°C. This fits into earlier observations of general reductions of temperature-induced expression plasticity in temperate compared to tropical populations collected from different locations across different continents (Levine et al. 2011; Zhao et al. 2015). More specifically, it supports the idea that temperature-sensitive expression of PcG target genes can be detrimental, and thus has been buffered in flies adapted to temperate climates in order to maintain more consistent expression levels across temperatures (Levine and Begun 2008). As an alternative explanation, since PcG proteins also function in regenerative processes (Grossniklaus and Paro 2014; Chou et al. 2019; Harris et al. 2020), might be that an upregulation of PcG target genes serves to control damages resulting from stress experienced at lower temperatures. Being better adapted to colder conditions (Hoffmann et al. 2003; Ayrinhac et al. 2004; Svetec et al. 2011; Pool et al. 2017), decreased temperature might cause less stress for temperate flies, and hence less upregulation of PcG target genes. An earlier study examining the expression of several PcG targets, however, observed a stronger reduction of PcG target gene overexpression at 15°C in warm-temperate flies from the French sample than in cold-temperate flies from a Swedish population sample (Voigt and Kost 2021). Overexpression at 15° was higher in Swedish compared to French flies, but still lower than in tropical Zambian flies. These differences in temperature-induced expression plasticity might result from the range in which temperatures vary in different climates. In warm-temperate climates the range encompasses cold and hot temperatures with greater regularity, whereas in cold-temperate and tropical climates the range is generally restricted to colder or hotter temperatures, respectively.

PcG and TrxG proteins regulate PcG target genes in an antagonistic manner, which includes establishing specific histone modifications characteristic of repressed and activated epigenetic states, respectively. The histone mark H3K27me3 is typical for PcG-mediated repression, while TrxG-typical H3K4me3 is associated with transcriptional activation (Kassis and Brown 2013; Simon and Kingston 2013; Steffen and Ringrose 2014; Entrevan et al. 2016; Giner-Laguarda and Vidal 2020; Kuroda et al. 2020). Both histone marks showed temperature-sensitive enrichment at PcG target genes in our study. Similar to PcG target gene expression, temperature-sensitive enrichment of H3K4me3 was more pronounced in females than in males, and more PcG targets harbored increased enrichment at 15°C than at 28°C in females. Furthermore, the temperature response of H3K4me3 enrichment was also significantly correlated to that of expression. That H3K4me3 enrichment can be increased at lower temperature resulting in higher expression at lower compared to higher temperatures was recently demonstrated for a one-gene locus (Gibert et al. 2016). Our study now gives further evidence for this on a genome-wide scale. Temperature-sensitive enrichment was less observed for H3K27me3. Temperature-sensitivity of H3K27me3 at PcG target sites, however, consisted in large parts of higher enrichment at 28°C. Thus, given that H3K27me3 is associated with transcriptional repression, temperature-sensitivities of both histone marks are consistent with more increased transcriptional activity at lower temperature, similar to what was observed for PcG target gene expression and to what has been classically found for PcG regulation (Fauvarque and Dura 1993; Chan et al. 1994; Zink and Paro 1995; Bantignies et al. 2003; Gibert et al. 2011; Voigt et al. 2015; Voigt et al. 2019; Voigt and Kost 2021). As for expression, the amount of temperature-sensitive enrichment of histone marks differed between temperate and tropical population samples, and was consistent with less overall transcriptional activation of PcG targets at lower temperature in temperate flies. For females, the proportion of PcG targets with higher H3K4me3 enrichment at 15°C was strongly reduced in temperate compared to tropical flies; whereas the reduction of H3K27me3 overenrichment at 28°C in temperate flies was stronger in males than females.

Both trans-regulatory and cis-regulatory factors might be responsible for the observed differences in temperature-sensitivity between temperate and tropical flies. Several genes encoding for products classified as TrxG proteins showed significant differences in their transcriptional response to temperature between temperate and tropical female flies, which might be related to the differences observed in temperature-sensitive H3K4me3 enrichment. The gene encoding for the PcG recruiter Grainyhead (Grh) also exhibited significant differences in its temperature response between temperate and tropical flies. Grh is known to interact with PREs as well as other PcG recruiter proteins such as Pleiohomeotic (Pho) and Pleiohomeotic-like (Phol) (Blastyák et al. 2006). DNA binding motifs for Pho and Phol were overrepresented and significantly differentiated between the French and Zambian populations at PcG target genes with population-specific temperature responses. Pho and Phol are part of the PcG complex PhoRC that functions in PcG recruitment and has a central role in PcG silencing (Brown et al. 1998; Wang et al. 2004; Klymenko et al. 2006; Kassis and Brown 2013; Frey et al. 2016). They are also subunits of the INO80 nucleosome remodeling complex (Klymenko et al. 2006), but also appear to have additional PcG-/INO80-independent functions in cell survival (Elizarev et al. 2021). Although their genome-wide binding distributions differ (Kahn et al. 2014), Pho and Phol appear to share partially redundant functions and DNA binding sites (Brown et al. 2003). Since their DNA binding motifs also share similarities, it cannot be ruled out that the observed motif differentiation might only pertain to one of the two factors. Pho was shown to strongly colocalize with proteins binding to insulator elements (Nègre et al. 2010). DNA binding motifs for such insulator proteins were also found to be overrepresented and significantly differentiated between the populations at PcG target genes with population-specific temperature responses. Insulator proteins are classically associated with demarcating the boundaries of chromatin domains in order to block the spreading of repressive chromatin and inhibit enhancer-promoter contacts (Moretti et al. 2020). Further studies revealed their roles in mediating long-range interactions between distant genomic sequences, and in the formation of the 3D chromatin structure (Moretti et al. 2020; Peterson et al. 2021). Moreover, the involvement of insulator proteins in PcG function becomes more and more apparent. They are involved in long-range spreading of H3K27me3 (Heurteau et al. 2020), interact with many of the PcG recruiter proteins (Chetverina et al. 2022a), and are often located next to PREs (Hagstrom et al. 1997; Mihaly et al. 1997; Gruzdeva et al. 2005; Kyrchanova et al. 2018) whose function they are able to enhance and even induce (Erokhin et al. 2021; Chetverina et al. 2022b). Binding sites for insulator proteins appear to evolve more swiftly than those of other transcription factors (Nègre et al. 2010). For example, binding of CTCF, the insulator protein with significant motif differentiation and overrepresentation across both sexes in our study, shows a highly dynamic evolution and can evolve adaptively (McDaniell et al. 2010; Ni et al. 2012). Point mutation can lead to changes in CTCF binding (Katainen et al. 2015), and such binding changes due to cis-regulatory changes have been associated with altered expression of adjacent genes in *Drosophila* (Ni et al. 2012). Interestingly, the strength of an insulator correlates to the density of bound insulator proteins, and furthermore, mutations in binding sites of insulator proteins can lead to reorganization of chromatin structure as well as gene regulatory changes (Van Bortle et al. 2014; Guo et al. 2015; Lupiáñez et al. 2015; Narendra et al. 2015). Since other transcription factors are involved in determining the transcriptional states of PcG targets, which are then epigenetically set by PcG/TrxG proteins (Schuettengruber and Cavalli 2009), we also looked for other factors exhibiting significant motif differentiation and overrepresentation at PcG targets with population-specific temperature responses in their expression. The Ecdysone receptor (EcR) was such a factor with significant motif differentiation and overrepresentation in both females and males. Interestingly, EcR is known to interact with many TrxG proteins (Badenhorst et al. 2005; Kirilly et al. 2011; Chauhan et al. 2012; Denton et al. 2013; Kreher et al. 2017; Shokri et al. 2019; Mazina et al. 2020), including those for which we observed population-specific temperature-sensitive expression (Osa & Mi-2), as well as PcG proteins, PcG recruiters (Hitrik et al. 2016; Gutierrez-Perez et al. 2019) and insulator proteins such as CTCF (Pascual-Garcia et al. 2017; Mazina et al. 2020). Recently, it has been shown that EcR can function together with the PcG protein Pc and the PcG recruiter Psq to alter 3D chromatin structure resulting in increased gene expression (Gutierrez-Perez et al. 2019). Altogether, this hints that differences in temperature-sensitive expression between temperate and tropical flies might involve structural changes of the 3D chromatin organization, however, further studies are needed to confirm this.

## Material and methods

### Fly lines and sample collection

Each population sample consisted of isofemale, inbred lines derived from natural populations from France and Zambia (Pool et al. 2012; Lack et al. 2016) (table 1). Both populations are well-studied and eight lines from each were randomly chosen for this study. All lines were reared in separate vials, on standard cornmeal-sucrose yeast medium with a light:dark cycle of 12:12 hours. In order to control for generational effects, parental flies of the experimental generation were reared at 21°C and transferred after hatching to the experimental temperatures of 15°C and 28° for mating and oviposition. The resulting progeny was then reared at the respective temperature under density-controlled conditions (50 larvae per vial). Seven-day old mated adult flies (males and females, separately) from each line were collected, flash-frozen in liquid nitrogen and stored at -80°C. Sex- and temperature-specific samples were collected for each population by evenly pool the eight lines of each population.

### RNA-seq analysis

RNA was extracted using the MasterPure Complete DNA and RNA Purification Kit (Lucigen). Samples consisted of pools of two individuals per line resulting in 16 flies per each sex-, temperature-, and population-specific sample. Three biological replicates were done for each sample. Library preparations were performed using NEBNext Ultra II RNA Library Prep Kit for Illumina. Over 60 million 2x100bp (PE) reads per sample were sequenced on an Illumina NovaSeq 6000 platform. Reads were mapped to the reference genome of *D. melanogaster* (Flybase Release 6.40; Gramates et al. 2022) with *STAR* (version v2.7.9a; Dobin et al. 2013). Read counts were then subjected to *edgeR* (version 3.30.3; Robinson et al. 2010) in order to identify differential expressed genes. Models were fitted using the generalized linear modelling (glm) approach and contrasts were examined with likelihood ratio tests. The Benjamini-Hochberg method was employed for multiple testing correction (Benjamini and Hochberg 1995). For each sex, contrasts were set up to assess temperature effects within each population (i.e., [France at 15°C – France at 28°C] and [Zambia at 15°C – Zambia at 28°C]), as well as interaction effects between population and temperature (i.e., [France at 15°C – France at 28°C] – [Zambia at 15°C – Zambia at 28°C]). All PcG target genes as given in Schwartz et al. (2010) were considered in this study. Schwartz et al. (2010) identified PcG targets based on the binding of PcG proteins and the histone modification H3K27me3. According to the current Flybase release 6.40 (Gramates et al. 2022), this corresponds to 818 PcG target genes in total. PcG target genes with significant expression differences were subjected to overrepresentation analyses using clusterProfiler (version 4.0.5; Wu et al. 2021) based on the genome annotation for *D. melanogaster* (Carlson 2021). Significant gene ontology (GO) terms of biological processes (BP) were derived by comparing the PcG target gene sets to a background sample consisting of all PcG target genes with observed expression. Multiple testing correction was done using the Benjamini-Hochberg method (Benjamini and Hochberg 1995).

### ChIP-seq analysis

Chromatin immunoprecipitation followed by high-throughput sequencing (ChIP-seq) was performed against the histone modifications trimethylation of histone 3 at lysine 4 (H3K4me3) and trimethylation of histone 3 at lysine 27 (H3K27me3). Native ChIP was employed using micrococcal nuclease (Mnase) digestion for chromatin fragmentation, which is considered the most-suited method for histone modifications (Kidder et al. 2011). Samples consisted of pools of four individuals per line resulting in 32 flies per each sex-, temperature-, and population-specific sample. Two biological replicates were done for each sample. Samples were placed in 1ml of buffer 1 containing 0.3 M sucrose, 30 mM KCl, 7.5 mM NaCl, 2.5 mM MgCl_2_, 0.05 mM EDTA, 0.1 mM PMSF, 0.5 mM DTT, 7.5 mM Tris-HCL pH 7.5, 5mM sodium butyrate, and Roche cOmplete Protease Inhibitor Cocktail tablets (1 tablet per 50 ml buffer). After adding 1 ml of lysis buffer (buffer 1 with 1% NP40), samples were homogenized on ice using a Dounce tissue grinder for 10 min. Samples were then overlaid on 8 ml of a buffer similar to buffer 1, except of a lower sucrose concentration of 1.2 M, in Corex centrifuge tubes and centrifuged for 20 min (8500 rpm at 4°C). The pellet was resuspended in 0.25 ml of Mnase digestion buffer (0.12 M sucrose, 0.2 mM PMSF, 4 mM MgCl_2_, 1 mM CaCl2, 0.05 Tris-HCl pH 7.5, 5 mM sodium butyrate). Chromatin digestion was then started by adding 15 U of Mnase (Thermo Scientific) to each sample. Digestion was stopped after 6 min at 37°C by adding 10 µl of 0.5 M EDTA and putting the samples on ice. After 10 min of centrifugation (13,000g at 4°C), samples were splitted into pellets and supernatants (S1). Supernatants were stored at -20°C, while each pellet was suspended into 100 µl dialysis buffer (200 µM EDTA, 200 µM PMSF, 1 mM Tris-HCl pH 7.5, 5mM sodium butyrate) and dialyzed against 50 ml dialysis buffer overnight at 4°C using Slide-A-Lyzer 3.5K MWCO MINI Dialysis Devices (Thermo Scientific). The dialyzed samples were then centrifuged for 10 min (13,000g at 4°C) and each supernatant was combined with the corresponding supernatant S1. In order to minimize unspecific backgrounds, samples were then centrifuged three times for 10 min and each time the supernatants were transferred into fresh tubes. Immunoprecipitation incubation buffer (150 mM NaCl, 5mM EDTA, 100 µM PMSF, 20 mM Tris-HCl pH 7.5, 20mM sodium butyrate) and antibodies against H3K4me3 (Merck/Millipore 04-745) or H3K27me3 (Merck/Millipore 07-449) were added to the samples. Half of the volume of each sample was incubated without antibodies (AB-control). After incubating overnight at 4°C, 50 µl of protein A-sepharose (Sigma) were added to each sample. Samples were incubated for 4 hours at 4°C, and then centrifuges for 10 min (11,660g at 4°C). Supernatants of AB-controls were stored at -20°C as “input” controls. Pellets were suspended in 10 ml washing buffer (75 mM NaCl, 10 mM EDTA, 50 mM Tris-HCl pH 7.5, 5 mM sodium butyrate), mixed gently for 10 min at 4°C, and after that centrifuged for 10 min (4000rpm at 4°C). This was repeated twice with 10 ml of the same washing buffer but increasing NaCl concentrations (125 mM and 175 mM, respectively). DNA of samples and input controls was then extracted with the QIAquick PCR purification kit (Qiagen). Samples and input controls were used for library preparations with NEBNext Ultra II DNA Library Prep Kit for Illumina. Over 25 million 2x100bp (PE) reads per sample were sequenced on an Illumina NovaSeq 6000 platform. Prior to mapping, raw FASTQ reads were trimmed and filtered to remove adapter sequences and low-quality reads (minimum base PHRED quality of 20 and minimum read length of 75bp) using *cutadapt* (version 2.4; Martin 2011). Reads were mapped to the reference genome of *D. melanogaster* (Flybase Release 6.40) with *bowtie2* (version 2.3.5.1; (Langmead and Salzberg 2012). Subsequent quality filtering included removing reads without proper pairing, unmapped reads and those with a mapping quality below 30 using *SAMtools* (version 1.9; Li et al. 2009). Based on the ENCODE blacklist (Amemiya et al. 2019), genomic regions that frequently produce noise and artifacts were also removed with *BEDTools* (version 2.28.0; Quinlan 2014). Peaks were called using MACS2 (version 2.2.7.1; Zhang et al. 2008) with the broad option employed for H3K27me3 samples and default option (narrow) for H3K4me3 samples. A consensus peak set across all samples was generated using DiffBind (versions 2.16.2 & 3.2.4; Ross-Innes et al. 2012), by which peaks were re-centered and trimmed based on their summits (points of greatest overlap of reads) to produce more standardized peak intervals and reduce the influence from nonsignificant background reads. Summits length was ±200bp for H3K4me3 and ±300bp for H3K27me3 peaks, and was based on the respective distribution of peak widths. Raw read counts of consensus peaks (minus read counts from input controls) were submitted to *edgeR* (version 3.30.3; Robinson et al. 2010) to identify differential enrichment in histone modifications at PcG target genes. Based on the highest correlation between gene expression and enrichment of histone modifications, all H3K4me3 peaks within ±0.5kB and all H3K27me3 peaks within ±2.5kB around transcription start sites (TSSs) of PcG target genes were considered in the analysis. Models were fitted using the generalized linear modelling (glm) approach and contrasts were examined with likelihood ratio tests. The Benjamini-Hochberg method was employed for multiple testing correction (Benjamini and Hochberg 1995). For each sex, contrasts were set up to assess temperature effects within each population (i.e., [France at 15°C – France at 28°C] and [Zambia at 15°C – Zambia at 28°C]), as well as interaction effects between temperature and population (i.e., [France at 15°C – France at 28°C] – [Zambia at 15°C – Zambia at 28°C]).

### Motif differentiation analysis

Sequence data of multiple genomes from the French (96 lines) and Zambian (197 lines) populations studied here were retrieved from the population genomic resource *Drosophila* Genome Nexus (DGN) (Lack et al. 2016). *F_ST_* (Weir and Cockerham 1984) per SNP was estimated in the euchromatic regions (i.e., regions of nonzero recombination) of the genome (Fiston-Lavier et al. 2010) along the major chromosome arms (X, 2L, 2R, 3L, and 3R) using *VCFtools* (version 0.1.17; Danecek et al. 2011). SNPs with less than ten lines without missing data in one or both populations were excluded from the analysis. The empirical distribution of *F_ST_* values was derived for each chromosome arm and SNPs with *F_ST_* values above the 99^th^ quantile were considered as outliers. Motif differentiation scores between the French and Zambian populations were calculated for fragments of ±20bp around outlier SNPs using *PWMenrich* (version 4.28.1; Stojnic and Diez 2022) without the background function. All position-weight matrices (PWMs) for *D. melanogaster* motifs included in *PWMenrich*, as well as the PWM for Dsp1, which was directly retrieved from JASPAR (Castro-Mondragon et al. 2022), were considered in the analysis. Redundant motifs were then removed based on motif consensus sequences (Table S35). Scores for each motif were first calculated for a background set of 10,000 (i.e., 2000 per chromosome arm) *F_ST_* outlier SNPs randomly drawn from gene regions (i.e., gene ± 2kb upstream and downstream of the gene) within the genome, and the 97.5^th^ quantiles of the resulting empirical distributions were considered as the threshold values of significant differentiation for each motif. Motifs with low motif differentiation were removed from the analysis. Motif differentiation scores between the two populations were then calculated for *F_ST_* outlier SNPs within the ± 2kb gene regions of PcG targets showing significant population x temperature interactions in their expression, and numbers of significant motif changes were counted based on the thresholds derived from the background set. For assessing which motif changes are over-represented at PcG targets with interaction effects in their expression, the number of significant motif changes for each motif in these PcG target regions was compared to that observed in PcG target regions with no significant temperature x population interactions in their expression and enrichment of histone modifications by employing Fisher’s exact tests and multiple testing correction according to Benjamini and Hochberg (Benjamini and Hochberg 1995).

## Data Availability

Fly lines are available upon request. Sequence data will be submitted to NCBI GenBank and accession numbers will be given here. The computational pipeline is given in the supplementary file 1.

## Acknowledgements

We are very grateful to the Klaus Reinhardt lab for providing the infrastructure and support that enabled our research, as well as Suzanne Eaton for her excellent input and support. She is deeply missed. This work was supported by the Deutsche Forschungsgemeinschaft (DFG) Research Unit grant VO 2380/1-1.

## Supplementary material – Figures

**Fig. S1.**
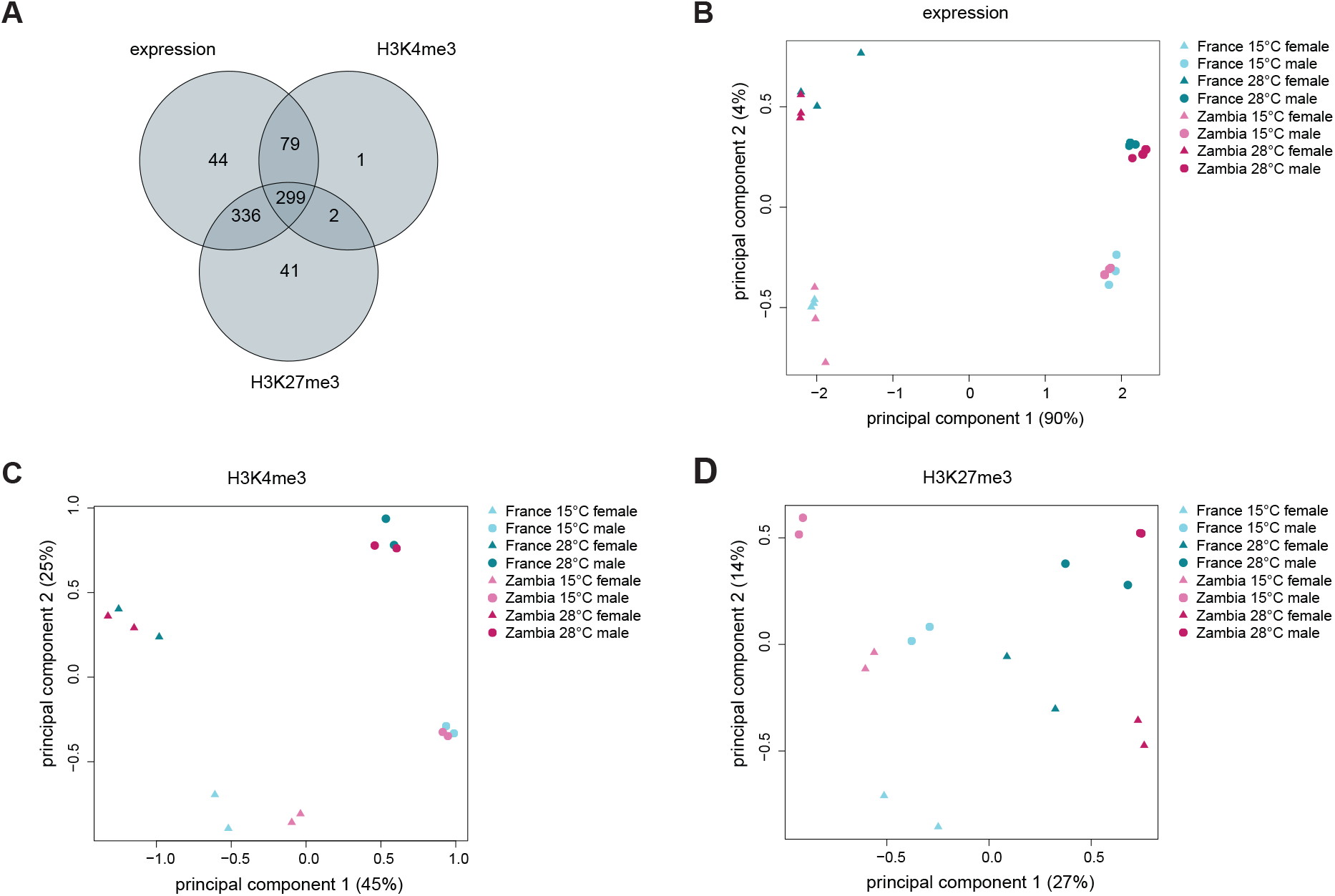
Differential expression and histone modification enrichment analyses of PcG targets in adult flies from temperate French and tropical Zambian populations. (A) Venn diagram showing numbers of PcG target genes with observed expression and enrichment of histone modifications H3K4me3 and H3K27me3 in present study, as well as their overlaps. (B) Principal component analyses on expression levels of PcG target genes, as well as on enrichment levels of (C) H3K4me3 and (D) H3K27me3 at PcG targets. The first and second principal components are shown, and the percentages of variation explained by each component are given in parentheses.

**Fig. S2.**
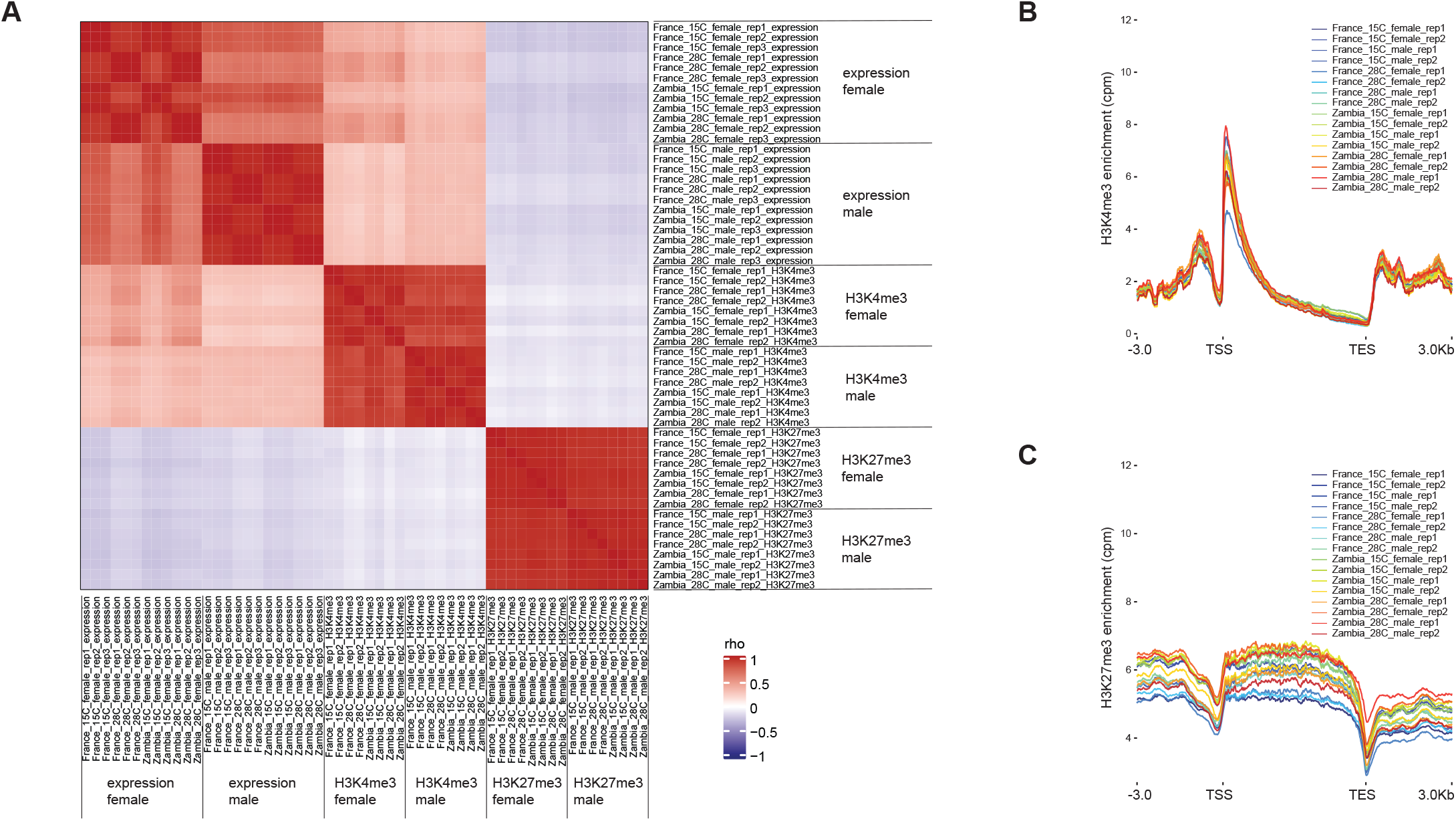
Quality control of ChIP-seq and RNA-seq data. (A) Spearman’s rank correlation coefficients (rho) among PcG target gene expression levels as well as associated H3K4me3 and H3K27me3 enrichment levels of all replicates. Profile plots of average (B) H3K4me3 and (C) H3K27me3 enrichment along PcG target gene regions.

**Fig. S3.**
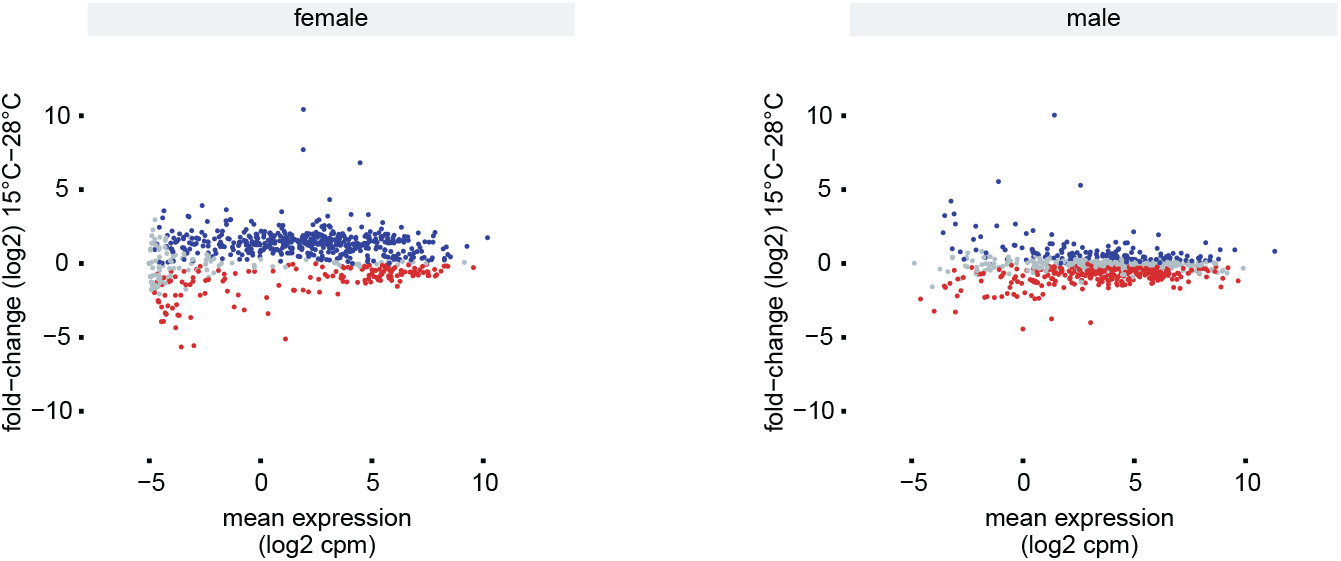
Expression levels of PcG target genes and corresponding temperature responses. Mean 15/28°C foldchanges per population sample and rearing temperature are plotted against mean expression levels. Blue and red dots correspond to significant overexpression at 15°C and at 28°C (FDR < 5%), respectively.

**Fig. S4.**
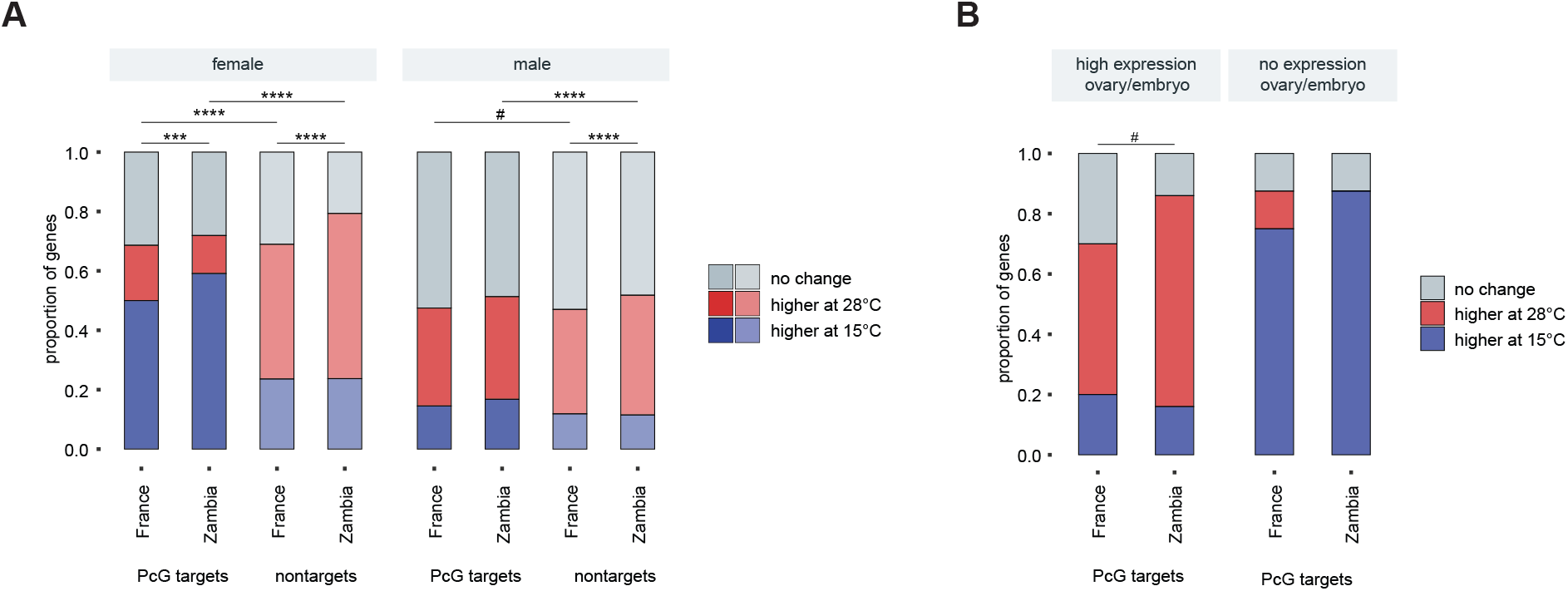
Proportions of PcG target genes with overexpression at 15°C (blue) and 28°C (red) (A) compared to genes not targeted by PcG proteins, as well as (B) of those PcG targets with high and no expression in ovaries and embryos.

**Fig. S5.**
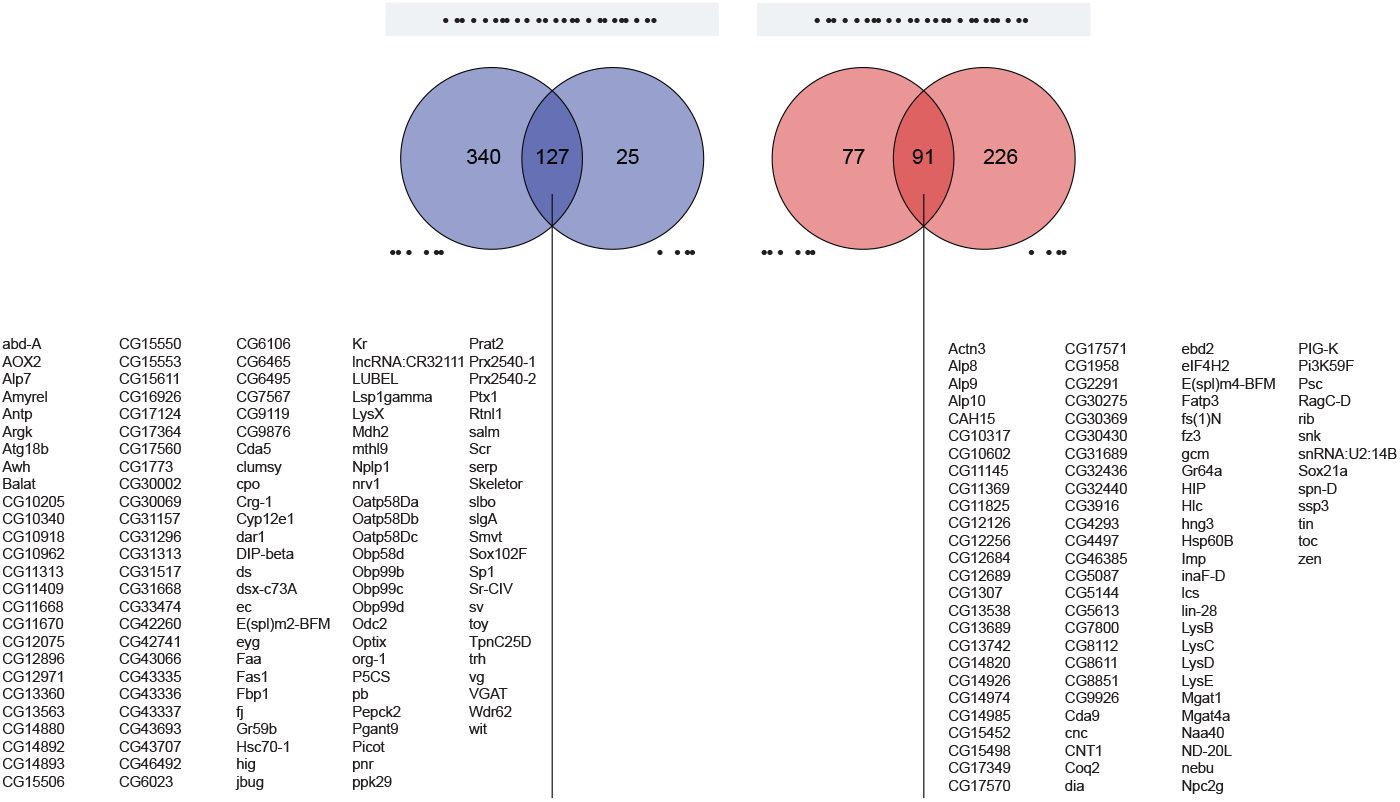
Female-male overlaps of PcG target genes with significant temperature-sensitive expression (FDR < 5%).

**Fig. S6.**
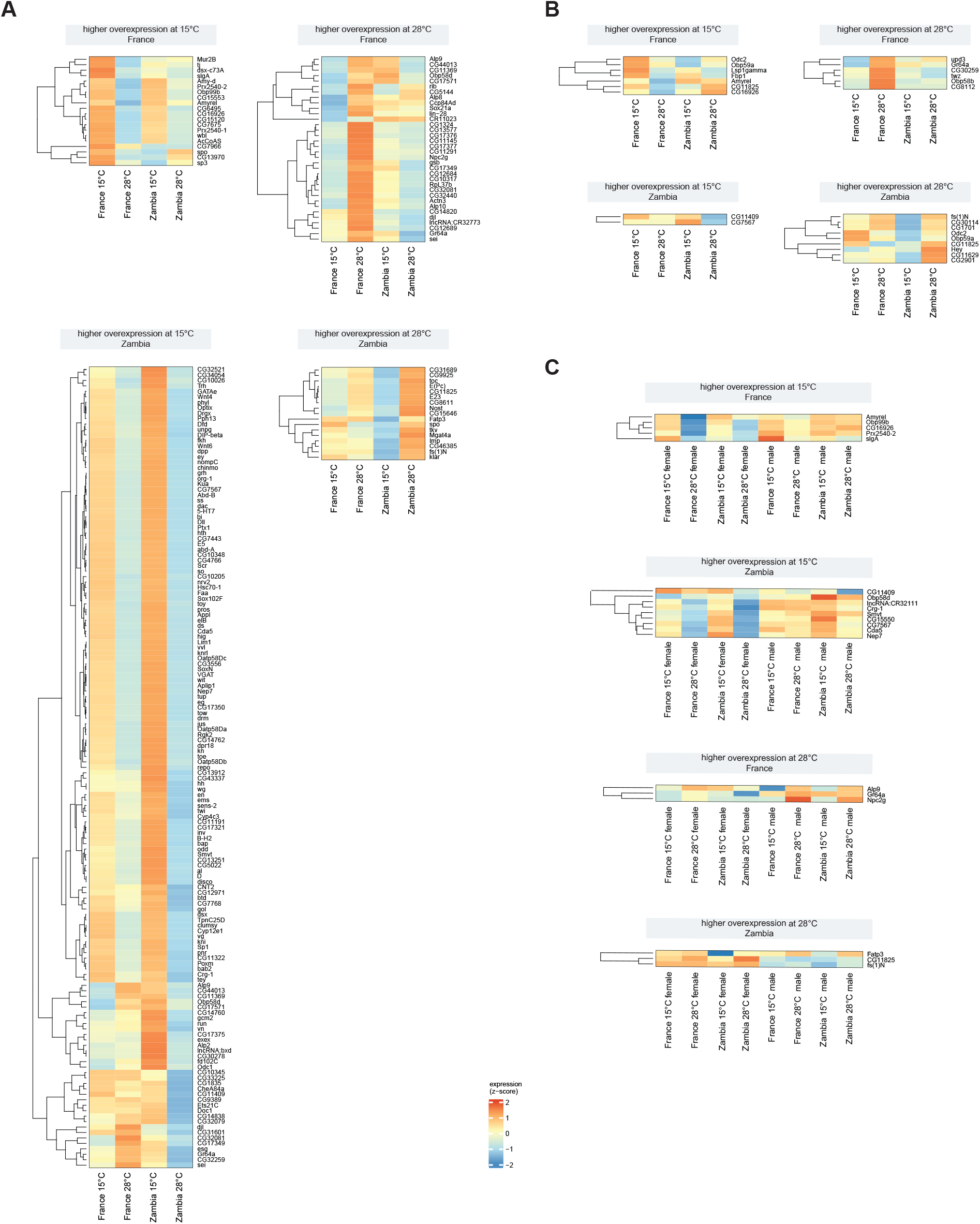
Expression levels across temperatures of PcG targets genes with significant population x temperature interaction effects in their expression (FDR < 5%). Gene-wise standardized expression levels (Z scores) are given.

**Fig. S7.**
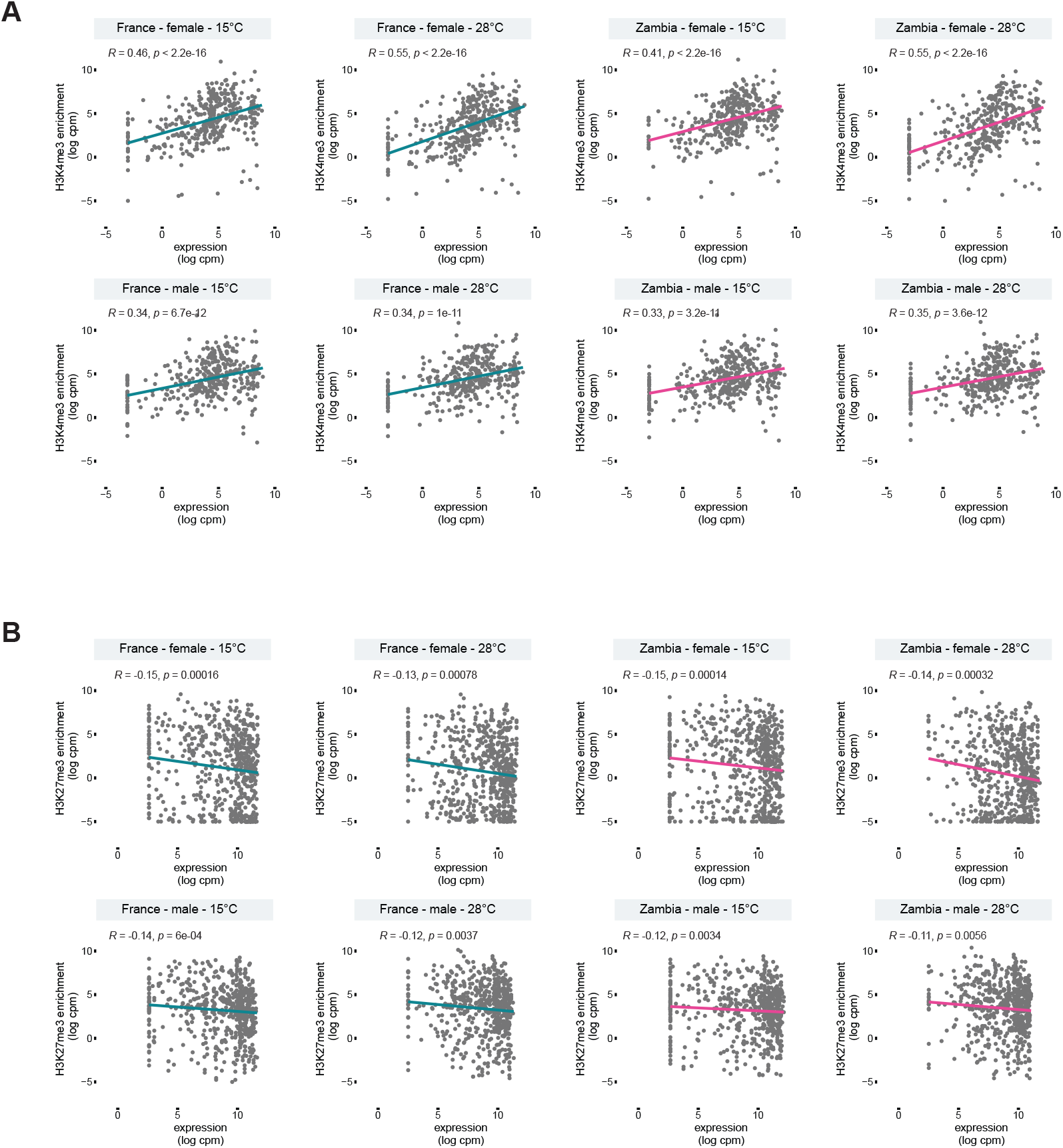
Correlation between PcG target gene expression and enrichment levels of (A) H3K4me3 and (B) H3K27me3. Spearman’s rank correlation coefficient (R) was estimated.

**Fig. S8.**
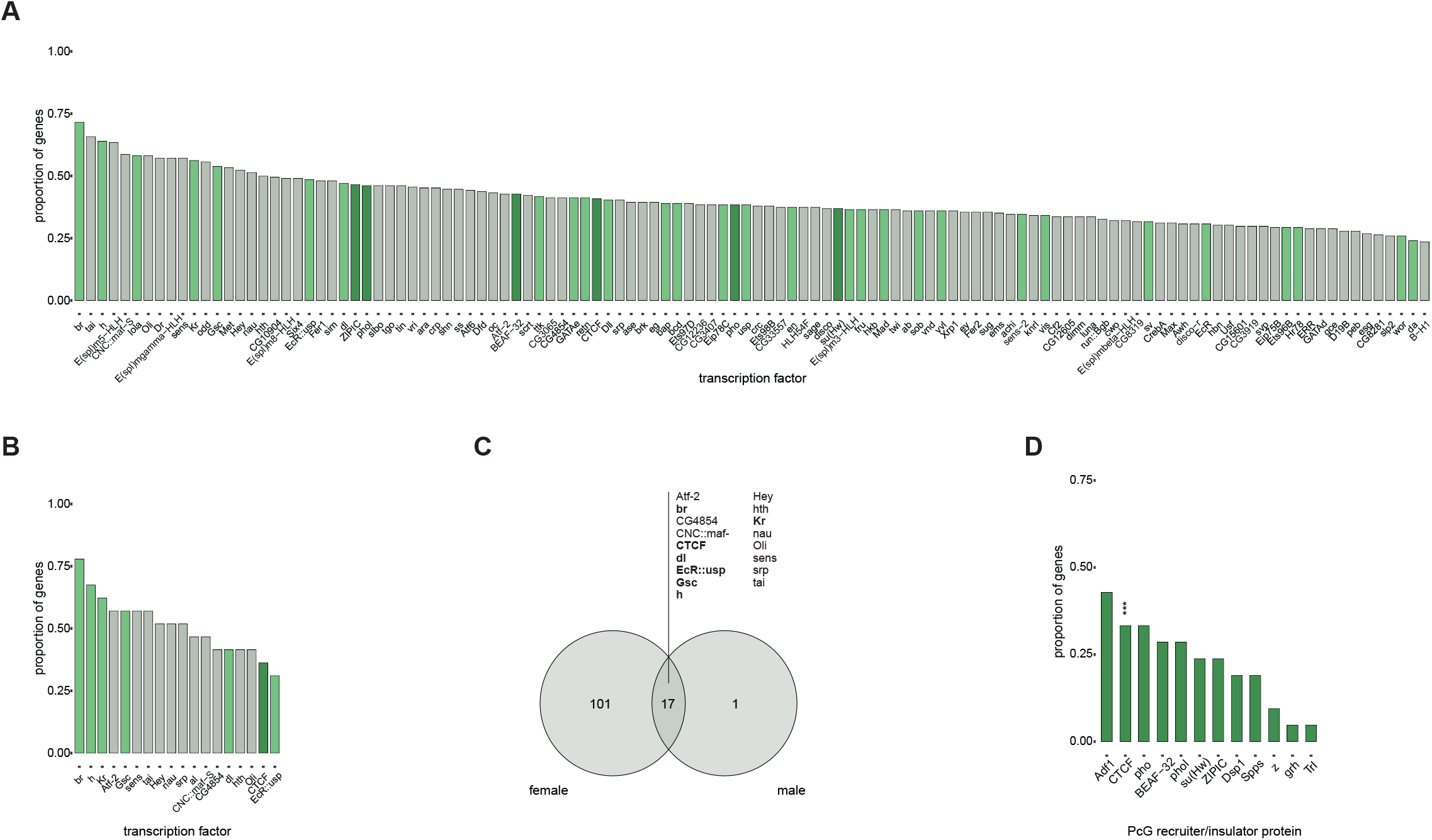
Candidates for cis-acting factors involved in differences in temperature-sensitive PcG target gene expression between temperate and tropical populations. Proportions of PcG target genes with significant population x temperature interactions in their expression that harbor significant differentiation of DNA binding motifs between the populations in their gene regions. Transcription factors with overrepresented motif differentiation (FDR < 5%) at PcG targets with significant interaction effects are given for (A) females and (B) males. Overrepresentation was assessed against the number of differentiated motifs found in gene regions of PcG targets with no significant population x temperature interactions effects in their expression. Dark green corresponds to PcG recruiter and insulator proteins, while light green to factors that were shown to physically interact with TrxG-/PcG-associated proteins (see also supplementary table S38) (C) Female-male overlap of transcription factors with significantly differentiated and overrepresented motifs. Factors with known interactions with PcG/TrxG-associated proteins are given in bold. (D) Proportions of PcG target genes with significant population x temperature interactions in their expression in males that harbor significant differentiation of DNA binding motifs of PcG recruiter and insulator proteins between the populations in their gene regions. Asterisks correspond to significant overrepresentation (FDR < 5%) of differentiated motifs at PcG target genes with significant interaction effects compared to those with none. *** *P* < 0.001.

## References

Amemiya HM, Kundaje A, Boyle AP. 2019. The ENCODE Blacklist: Identification of Problematic Regions of the Genome. Sci Rep 9:9354.

Arguello JR, Laurent S, Clark AG. 2019. Demographic History of the Human Commensal Drosophila melanogaster. Genome Biology and Evolution 11:844–854.

Ayrinhac A, Debat V, Gibert P, Kister A-G, Legout H, Moreteau B, Vergilino R, David JR. 2004. Cold adaptation in geographical populations of Drosophila melanogaster: phenotypic plasticity is more important than genetic variability. Functional Ecology 18:700–706.

Badenhorst P, Xiao H, Cherbas L, Kwon SY, Voas M, Rebay I, Cherbas P, Wu C. 2005. The Drosophila nucleosome remodeling factor NURF is required for Ecdysteroid signaling and metamorphosis. Genes Dev 19:2540–2545.

Bantignies F, Grimaud C, Lavrov S, Gabut M, Cavalli G. 2003. Inheritance of Polycomb-dependent chromosomal interactions in Drosophila. Genes Dev. 17:2406–2420.

Benjamini Y, Hochberg Y. 1995. Controlling the False Discovery Rate: A Practical and Powerful Approach to Multiple Testing. Journal of the Royal Statistical Society. Series B (Methodological) 57:289–300.

Blastyák A, Mishra RK, Karch F, Gyurkovics H. 2006. Efficient and Specific Targeting of Polycomb Group Proteins Requires Cooperative Interaction between Grainyhead and Pleiohomeotic. Molecular and Cellular Biology 26:1434–1444.

Bonchuk A, Denisov S, Georgiev P, Maksimenko O. 2011. Drosophila BTB/POZ Domains of “ttk Group” Can Form Multimers and Selectively Interact with Each Other. Journal of Molecular Biology 412:423–436.

Bornelöv S, Reynolds N, Xenophontos M, Gharbi S, Johnstone E, Floyd R, Ralser M, Signolet J, Loos R, Dietmann S, et al. 2018. The Nucleosome Remodeling and Deacetylation Complex Modulates Chromatin Structure at Sites of Active Transcription to Fine-Tune Gene Expression. Mol Cell 71:56–72.e4.

Boulétreau-Merle J, Terrier O. 1986. Adaptive diversity in genetic control of egg-laying behavior in Drosophila melanogaster. International Journal of Invertebrate Reproduction and Development 9:113–124.

Bray SJ, Kafatos FC. 1991. Developmental function of Elf-1: an essential transcription factor during embryogenesis in Drosophila. Genes Dev. 5:1672–1683.

Brown JL, Fritsch C, Mueller J, Kassis JA. 2003. The Drosophila pho-like gene encodes a YY1-related DNA binding protein that is redundant with pleiohomeotic in homeotic gene silencing. Development 130:285–294.

Brown JL, Mucci D, Whiteley M, Dirksen M-L, Kassis JA. 1998. The Drosophila Polycomb Group Gene pleiohomeotic Encodes a DNA Binding Protein with Homology to the Transcription Factor YY1. Molecular Cell 1:1057–1064.

Brown JL, Sun M, Kassis JA. 2018. Global changes of H3K27me3 domains and Polycomb group protein distribution in the absence of recruiters Spps or Pho. Proceedings of the National Academy of Sciences 115:E1839–E1848.

Carlson M. 2021. org.Dm.eg.db: Genome wide annotation for Fly. R package version 3.13.0. Available from: http://bioconductor.org/packages/org.Dm.eg.db/

Castro-Mondragon JA, Riudavets-Puig R, Rauluseviciute I, Berhanu Lemma R, Turchi L, Blanc-Mathieu R, Lucas J, Boddie P, Khan A, Manosalva Pérez N, et al. 2022. JASPAR 2022: the 9th release of the open-access database of transcription factor binding profiles. Nucleic Acids Research 50:D165–D173.

Cavalli G, Heard E. 2019. Advances in epigenetics link genetics to the environment and disease. Nature 571:489–499.

Chan C s., Rastelli L, Pirrotta V. 1994. A Polycomb response element in the Ubx gene that determines an epigenetically inherited state of repression. The EMBO Journal 13:2553–2564.

Chauhan C, Zraly CB, Parilla M, Diaz MO, Dingwall AK. 2012. Histone recognition and nuclear receptor co-activator functions of Drosophila Cara Mitad, a homolog of the N-terminal portion of mammalian MLL2 and MLL3. Development 139:1997–2008.

Chetverina D, Vorobyeva NE, Mazina MY, Fab LV, Lomaev D, Golovnina A, Mogila V, Georgiev P, Ziganshin RH, Erokhin M. 2022. Comparative interactome analysis of the PRE DNA-binding factors: purification of the Combgap-, Zeste-, Psq-, and Adf 1-associated proteins. Cell Mol Life Sci 79:353.

Chetverina DA, Gorbenko FV, Lomaev DV, Georgiev PG, Erokhin MM. 2022. Recruitment to Chromatin of (GA)n-Associated Factors GAF and Psq in the Transgenic Model System Depends on the Presence of Architectural Protein Binding Sites. Dokl Biochem Biophys 506:210–214.

Chetverina DA, Lomaev DV, Erokhin MM. 2020. Polycomb and Trithorax Group Proteins: The Long Road from Mutations in Drosophila to Use in Medicine. Acta Naturae 12:66–85.

Cheutin T, Cavalli G. 2012. Progressive Polycomb Assembly on H3K27me3 Compartments Generates Polycomb Bodies with Developmentally Regulated Motion. PLOS Genetics 8:e1002465.

Chou J, Ferris AC, Chen T, Seok R, Yoon D, Suzuki Y. 2019. Roles of Polycomb group proteins Enhancer of zeste (E(z)) and Polycomb (Pc) during metamorphosis and larval leg regeneration in the flour beetle Tribolium castaneum. Developmental Biology 450:34–46.

Classen A-K, Bunker BD, Harvey KF, Vaccari T, Bilder D. 2009. A tumor suppressor activity of Drosophila Polycomb genes mediated by JAK-STAT signaling. Nat Genet 41:1150–1155.

Comet I, Schuettengruber B, Sexton T, Cavalli G. 2011. A chromatin insulator driving three-dimensional Polycomb response element (PRE) contacts and Polycomb association with the chromatin fiber. PNAS 108:2294–2299.

Danecek P, Auton A, Abecasis G, Albers CA, Banks E, DePristo MA, Handsaker RE, Lunter G, Marth GT, Sherry ST, et al. 2011. The variant call format and VCFtools. Bioinformatics 27:2156–2158.

David JR, Capy P. 1988. Genetic variation of Drosophila melanogaster natural populations. Trends in Genetics 4:106–111.

Decoville M, Giraud-Panis MJ, Mosrin-Huaman C, Leng M, Locker D. 2000. HMG boxes of DSP1 protein interact with the Rel homology domain of transcription factors. Nucleic Acids Research 28:454– 462.

Delpuech J-M, Moreteau B, Chiche J, Pla E, Vouidibio J, David JR. 1995. Phenotypic Plasticity and Reaction Norms in Temperate and Tropical Populations of Drosophila Melanogaster: Ovarian Size and Developmental Temperature. Evolution 49:670–675.

Denton D, Aung-Htut MT, Lorensuhewa N, Nicolson S, Zhu W, Mills K, Cakouros D, Bergmann A, Kumar S. 2013. UTX coordinates steroid hormone-mediated autophagy and cell death. Nat Commun 4:2916.

Detienne G, De Haes W, Mergan L, Edwards SL, Temmerman L, Van Bael S. 2018. Beyond ROS clearance: Peroxiredoxins in stress signaling and aging. Ageing Research Reviews 44:33–48.

Dobin A, Davis CA, Schlesinger F, Drenkow J, Zaleski C, Jha S, Batut P, Chaisson M, Gingeras TR. 2013. STAR: ultrafast universal RNA-seq aligner. Bioinformatics 29:15–21.

Elizarev P, Finkl K, Müller J. 2021. Distinct requirements for Pho, Sfmbt, and Ino 80 for cell survival in Drosophila. Genetics 219:iyab096.

Enderle D, Beisel C, Stadler MB, Gerstung M, Athri P, Paro R. 2011. Polycomb preferentially targets stalled promoters of coding and noncoding transcripts. Genome Res. 21:216–226.

Entrevan M, Schuettengruber B, Cavalli G. 2016. Regulation of Genome Architecture and Function by Polycomb Proteins. Trends in Cell Biology 26:511–525.

Erokhin M, Gorbenko F, Lomaev D, Mazina MY, Mikhailova A, Garaev AK, Parshikov A, Vorobyeva NE, Georgiev P, Schedl P, et al. 2021. Boundaries potentiate polycomb response element-mediated silencing. BMC Biology 19:113.

Fauvarque MO, Dura JM. 1993. polyhomeotic regulatory sequences induce developmental regulator-dependent variegation and targeted P-element insertions in Drosophila. Genes Dev. 7:1508– 1520.

Fiston-Lavier A-S, Singh ND, Lipatov M, Petrov DA. 2010. Drosophila melanogaster recombination rate calculator. Gene 463:18–20.

Frey F, Sheahan T, Finkl K, Stoehr G, Mann M, Benda C, Müller J. 2016. Molecular basis of PRC1 targeting to Polycomb response elements by PhoRC. Genes Dev. 30:1116–1127.

Fu D, Wen Y, Ma J. 2004. The Co-activator CREB-binding Protein Participates in Enhancer-dependent Activities of Bicoid *. Journal of Biological Chemistry 279:48725–48733.

Gibert J-M, Karch F, Schlötterer C. 2011. Segregating Variation in the Polycomb Group Gene cramped Alters the Effect of Temperature on Multiple Traits. PLOS Genetics 7:e1001280.

Gibert J-M, Mouchel-Vielh E, Castro SD, Peronnet F. 2016. Phenotypic Plasticity through Transcriptional Regulation of the Evolutionary Hotspot Gene tan in Drosophila melanogaster. PLOS Genetics 12:e1006218.

Gibert J-M, Peronnet F. 2021. The Paramount Role of Drosophila melanogaster in the Study of Epigenetics: From Simple Phenotypes to Molecular Dissection and Higher-Order Genome Organization. Insects 12:884.

Giner-Laguarda N, Vidal M. 2020. Functions of Polycomb Proteins on Active Targets. Epigenomes 4:17.

González J, Macpherson JM, Petrov DA. 2009. A recent adaptive transposable element insertion near highly conserved developmental loci in Drosophila melanogaster. Mol Biol Evol 26:1949–1961.

Gramates LS, Agapite J, Attrill H, Calvi BR, Crosby MA, dos Santos G, Goodman JL, Goutte-Gattat D, Jenkins VK, Kaufman T, et al. 2022. FlyBase: a guided tour of highlighted features. Genetics 220:iyac035.

Grossniklaus U, Paro R. 2014. Transcriptional Silencing by Polycomb-Group Proteins. Cold Spring Harb Perspect Biol 6:a019331.

Gruzdeva N, Kyrchanova O, Parshikov A, Kullyev A, Georgiev P. 2005. The Mcp Element from the bithorax Complex Contains an Insulator That Is Capable of Pairwise Interactions and Can Facilitate Enhancer-Promoter Communication. Mol Cell Biol 25:3682–3689.

Guio L, Barrón MG, González J. 2014. The transposable element Bari-Jheh mediates oxidative stress response in Drosophila. Mol Ecol 23:2020–2030.

Guo Y, Xu Q, Canzio D, Shou J, Li J, Gorkin DU, Jung I, Wu H, Zhai Y, Tang Y, et al. 2015. CRISPR Inversion of CTCF Sites Alters Genome Topology and Enhancer/Promoter Function. Cell 162:900–910.

Guruharsha KG, Rual J-F, Zhai B, Mintseris J, Vaidya P, Vaidya N, Beekman C, Wong C, Rhee DY, Cenaj O, et al. 2011. A Protein Complex Network of Drosophila melanogaster. Cell 147:690–703.

Gutierrez-Perez I, Rowley MJ, Lyu X, Valadez-Graham V, Vallejo DM, Ballesta-Illan E, Lopez-Atalaya JP, Kremsky I, Caparros E, Corces VG, et al. 2019. Ecdysone-Induced 3D Chromatin Reorganization Involves Active Enhancers Bound by Pipsqueak and Polycomb. Cell Rep 28:2715–2727.e5.

Hagstrom K, Muller M, Schedl P. 1997. A Polycomb and Gaga Dependent Silencer Adjoins the Fab-7 Boundary in the Drosophila Bithorax Complex. Genetics 146:1365–1380.

Harris RE, Stinchfield MJ, Nystrom SL, McKay DJ, Hariharan IK. 2020. Damage-responsive, maturity-silenced enhancers regulate multiple genes that direct regeneration in Drosophila. Bellen HJ, Banerjee U, editors. eLife 9:e58305.

Heurteau A, Perrois C, Depierre D, Fosseprez O, Humbert J, Schaak S, Cuvier O. 2020. Insulator-based loops mediate the spreading of H3K27me3 over distant micro-domains repressing euchromatin genes. Genome Biology 21:193.

Hitrik A, Popliker M, Gancz D, Mukamel Z, Lifshitz A, Schwartzman O, Tanay A, Gilboa L. 2016. Combgap Promotes Ovarian Niche Development and Chromatin Association of EcR-Binding Regions in BR-C. PLOS Genetics 12:e1006330.

Hoffmann AA, Sørensen JG, Loeschcke V. 2003. Adaptation of Drosophila to temperature extremes: bringing together quantitative and molecular approaches. Journal of Thermal Biology 28:175– 216.

Holmqvist P-H, Boija A, Philip P, Crona F, Stenberg P, Mannervik M. 2012. Preferential Genome Targeting of the CBP Co-Activator by Rel and Smad Proteins in Early Drosophila melanogaster Embryos. PLOS Genetics 8:e1002769.

Horváth B, Kalinka AT. 2018. The genetics of egg retention and fertilization success in Drosophila: One step closer to understanding the transition from facultative to obligate viviparity. Evolution 72:318–336.

Hsiao Y-L, Chen Y-J, Chang Y-J, Yeh H-F, Huang Y-C, Pi H. 2014. Proneural proteins Achaete and Scute associate with nuclear actin to promote formation of external sensory organs. Journal of Cell Science 127:182–190.

Hu Y, Perrimon N, Vidal M, Celniker S. 2018. A Comprehensive Drosophila melanogaster Transcription Factor Interactome: Cell Reports. Available from: https://www.cell.com/cell-reports/fulltext/S2211-1247(19)30401-2?_returnURL=https%3A%2F%2Flinkinghub.elsevier.com%2Fretrieve%2Fpii%2FS2211124719304012%3Fshowall%3Dtrue

Huang Y, Agrawal AF. 2016. Experimental Evolution of Gene Expression and Plasticity in Alternative Selective Regimes. PLOS Genetics 12:e1006336.

Hur M-W, Laney JD, Jeon S-H, Ali J, Biggin MD. 2002. Zeste maintains repression of Ubx transgenes: support for a new model of Polycomb repression. Development 129:1339–1343.

Ito H, Sato K, Koganezawa M, Ote M, Matsumoto K, Hama C, Yamamoto D. 2012. Fruitless Recruits Two Antagonistic Chromatin Factors to Establish Single-Neuron Sexual Dimorphism. Cell 149:1327– 1338.

Kahn TG, Stenberg P, Pirrotta V, Schwartz YB. 2014. Combinatorial Interactions Are Required for the Efficient Recruitment of Pho Repressive Complex (PhoRC) to Polycomb Response Elements. PLOS Genetics 10:e1004495.

Kapopoulou A, Kapun M, Pieper B, Pavlidis P, Wilches R, Duchen P, Stephan W, Laurent S. 2020. Demographic analyses of a new sample of haploid genomes from a Swedish population of Drosophila melanogaster. Scientific Reports 10:22415.

Kassis JA, Brown JL. 2013. Chapter Three – Polycomb Group Response Elements in Drosophila and Vertebrates. In: Friedmann T, Dunlap JC, Goodwin SF, editors. Advances in Genetics. Vol. 81. Academic Press. p. 83–118. Available from: http://www.sciencedirect.com/science/article/pii/B9780124076778000038

Katainen R, Dave K, Pitkänen E, Palin K, Kivioja T, Välimäki N, Gylfe AE, Ristolainen H, Hänninen UA, Cajuso T, et al. 2015. CTCF/cohesin-binding sites are frequently mutated in cancer. Nat Genet 47:818–821.

Khalisova KY, Osadchiy IS, Georgiev PG, Maksimenko OG. 2021. TTK Isoforms Interact with Two Regions of the Mep-1 Protein of Drosophila melanogaster. Dokl Biochem Biophys 498:177–179.

Kidder BL, Hu G, Zhao K. 2011. ChIP-Seq: technical considerations for obtaining high-quality data. Nat Immunol 12:918–922.

Kirilly D, Wong JJL, Lim EKH, Wang Y, Zhang H, Wang C, Liao Q, Wang Haifeng, Liou Y-C, Wang Hongyan, et al. 2011. Intrinsic Epigenetic Factors Cooperate with the Steroid Hormone Ecdysone to Govern Dendrite Pruning in Drosophila. Neuron 72:86–100.

Klymenko T, Papp B, Fischle W, Köcher T, Schelder M, Fritsch C, Wild B, Wilm M, Müller J. 2006. A Polycomb group protein complex with sequence-specific DNA-binding and selective methyl-lysine-binding activities. Genes Dev. 20:1110–1122.

Kreher J, Kovač K, Bouazoune K, Mačinković I, Ernst AL, Engelen E, Pahl R, Finkernagel F, Murawska M, Ullah I, et al. 2017. EcR recruits dMi-2 and increases efficiency of dMi-2-mediated remodelling to constrain transcription of hormone-regulated genes. Nat Commun 8:14806.

Kuroda MI, Kang H, De S, Kassis JA. 2020. Dynamic Competition of Polycomb and Trithorax in Transcriptional Programming. Annu Rev Biochem 89:235–253.

Kyrchanova O, Kurbidaeva A, Sabirov M, Postika N, Wolle D, Aoki T, Maksimenko O, Mogila V, Schedl P, Georgiev P. 2018. The bithorax complex iab-7 Polycomb response element has a novel role in the functioning of the Fab-7 chromatin boundary. PLOS Genetics 14:e1007442.

Lachaise D, Silvain J-F. 2004. How two Afrotropical endemics made two cosmopolitan human commensals: the Drosophila melanogaster-D. simulans palaeogeographic riddle. In: Capy P, Gibert P, Boussy I, editors. Drosophila melanogaster, Drosophila simulans: So Similar, So Different. Contemporary Issues in Genetics and Evolution. Dordrecht: Springer Netherlands. p. 17–39. Available from: https://doi.org/10.1007/978-94-007-0965-2_2

Lack JB, Lange JD, Tang AD, Corbett-Detig RB, Pool JE. 2016. A Thousand Fly Genomes: An Expanded Drosophila Genome Nexus. Mol Biol Evol 33:3308–3313.

Langmead B, Salzberg SL. 2012. Fast gapped-read alignment with Bowtie 2. Nat Methods 9:357–359.

Laurent SJY, Werzner A, Excoffier L, Stephan W. 2011. Approximate Bayesian Analysis of Drosophila melanogaster Polymorphism Data Reveals a Recent Colonization of Southeast Asia. Mol Biol Evol 28:2041–2051.

Lehming N, Thanos D, Brickman JM, Ma J, Maniatis T, Ptashne M. 1994. An HMG-like protein that can switch a transcriptional activator to a repressor. Nature 371:175–179.

Levine MT, Begun DJ. 2008. Evidence of Spatially Varying Selection Acting on Four Chromatin-Remodeling Loci in Drosophila melanogaster. Genetics 179:475–485.

Levine MT, Eckert ML, Begun DJ. 2011. Whole-Genome Expression Plasticity across Tropical and Temperate Drosophila melanogaster Populations from Eastern Australia. Mol Biol Evol 28:249– 256.

Li H, Handsaker B, Wysoker A, Fennell T, Ruan J, Homer N, Marth G, Abecasis G, Durbin R, 1000 Genome Project Data Processing Subgroup. 2009. The Sequence Alignment/Map format and SAMtools. Bioinformatics 25:2078–2079.

Li H-B, Müller M, Bahechar IA, Kyrchanova O, Ohno K, Georgiev P, Pirrotta V. 2011. Insulators, Not Polycomb Response Elements, Are Required for Long-Range Interactions between Polycomb Targets in Drosophila melanogaster. Molecular and Cellular Biology 31:616–625.

Li H-B, Ohno K, Gui H, Pirrotta V. 2013. Insulators Target Active Genes to Transcription Factories and Polycomb-Repressed Genes to Polycomb Bodies. PLOS Genetics 9:e1003436.

Lilja T, Aihara H, Stabell M, Nibu Y, Mannervik M. 2007. The acetyltransferase activity of Drosophila CBP is dispensable for regulation of the Dpp pathway in the early embryo. Developmental Biology 305:650–658.

Lomaev D, Mikhailova A, Erokhin M, Shaposhnikov AV, Moresco JJ, Blokhina T, Wolle D, Aoki T, Ryabykh V, Iii JRY, et al. 2017. The GAGA factor regulatory network: Identification of GAGA factor associated proteins. PLOS ONE 12:e0173602.

Loubiere V, Delest A, Thomas A, Bonev B, Schuettengruber B, Sati S, Martinez A-M, Cavalli G. 2016. Coordinate redeployment of PRC1 proteins suppresses tumor formation during Drosophila development. Nature Genetics 48:1436–1442.

Luo L, Siah CK, Cai Y. 2017. Engrailed acts with Nejire to control decapentaplegic expression in the Drosophila ovarian stem cell niche. Development 144:3224–3231.

Lupiáñez DG, Kraft K, Heinrich V, Krawitz P, Brancati F, Klopocki E, Horn D, Kayserili H, Opitz JM, Laxova R, et al. 2015. Disruptions of Topological Chromatin Domains Cause Pathogenic Rewiring of Gene-Enhancer Interactions. Cell 161:1012–1025.

Lv X, Han Z, Chen H, Yang B, Yang X, Xia Y, Pan C, Fu L, Zhang S, Han H, et al. 2016. A positive role for polycomb in transcriptional regulation via H4K20me1. Cell Res 26:529–542.

Mallard F, Nolte V, Schlötterer C. 2020. The Evolution of Phenotypic Plasticity in Response to Temperature Stress. Genome Biology and Evolution 12:2429–2440.

Martin M. 2011. Cutadapt removes adapter sequences from high-throughput sequencing reads. EMBnet.journal 17:10–12.

Martinez A-M, Schuettengruber B, Sakr S, Janic A, Gonzalez C, Cavalli G. 2009. Polyhomeotic has a tumor suppressor activity mediated by repression of Notch signaling. Nat Genet 41:1076– 1082.

Mazina MY, Ziganshin RH, Magnitov MD, Golovnin AK, Vorobyeva NE. 2020. Proximity-dependent biotin labelling reveals CP190 as an EcR/Usp molecular partner. Sci Rep 10:4793.

McDaniell R, Lee B-K, Song L, Liu Z, Boyle AP, Erdos MR, Scott LJ, Morken MA, Kucera KS, Battenhouse A, et al. 2010. Heritable individual-specific and allele-specific chromatin signatures in humans. Science 328:235–239.

Mihaly J, Hogga I, Gausz J, Gyurkovics H, Karch F. 1997. In situ dissection of the Fab-7 region of the bithorax complex into a chromatin domain boundary and a Polycomb-response element. Development 124:1809–1820.

Moretti C, Stévant I, Ghavi-Helm Y. 2020. 3D genome organisation in Drosophila. Briefings in Functional Genomics 19:92–100.

Narendra V, Rocha PP, An D, Raviram R, Skok JA, Mazzoni EO, Reinberg D. 2015. CTCF establishes discrete functional chromatin domains at the Hox clusters during differentiation. Science 347:1017–1021.

Nègre N, Brown CD, Shah PK, Kheradpour P, Morrison CA, Henikoff JG, Feng X, Ahmad K, Russell S, White RAH, et al. 2010. A Comprehensive Map of Insulator Elements for the Drosophila Genome. PLOS Genetics 6:e1000814.

Ni X, Zhang YE, Nègre N, Chen S, Long M, White KP. 2012. Adaptive Evolution and the Birth of CTCF Binding Sites in the Drosophila Genome. PLOS Biology 10:e1001420.

Pagans S, Ortiz-Lombardía M, Espinás ML, Bernués J, Azorín F. 2002. The Drosophila transcription factor tramtrack (TTK) interacts with Trithorax-like (GAGA) and represses GAGA-mediated activation. Nucleic Acids Research 30:4406–4413.

Pagans S, Piñeyro D, Kosoy A, Bernués J, Azorín F. 2004. Repression by TTK69 of GAGA-mediated Activation Occurs in the Absence of TTK69 Binding to DNA and Solely Requires the Contribution of the POZ/BTB Domain of TTK69 *. Journal of Biological Chemistry 279:9725–9732.

Pascual-Garcia P, Debo B, Aleman JR, Talamas JA, Lan Y, Nguyen NH, Won KJ, Capelson M. 2017. Metazoan Nuclear Pores Provide a Scaffold for Poised Genes and Mediate Induced Enhancer-Promoter Contacts. Molecular Cell 66:63–76.e6.

Petavy G, David JR, Gibert P, Moreteau B. 2001. Viability and rate of development at different temperatures in Drosophila: a comparison of constant and alternating thermal regimes. Journal of Thermal Biology 26:29–39.

Peterson SC, Samuelson KB, Hanlon SL. 2021. Multi-Scale Organization of the Drosophila melanogaster Genome. Genes 12:817.

Pherson M, Misulovin Z, Gause M, Mihindukulasuriya K, Swain A, Dorsett D. 2017. Polycomb repressive complex 1 modifies transcription of active genes. Science Advances 3:e1700944.

Piunti A, Shilatifard A. 2016. Epigenetic balance of gene expression by Polycomb and COMPASS families. Science 352:aad9780.

Pool JE, Braun DT, Lack JB. 2017. Parallel Evolution of Cold Tolerance within Drosophila melanogaster. Mol Biol Evol 34:349–360.

Pool JE, Corbett-Detig RB, Sugino RP, Stevens KA, Cardeno CM, Crepeau MW, Duchen P, Emerson JJ, Saelao P, Begun DJ, et al. 2012. Population Genomics of Sub-Saharan Drosophila melanogaster: African Diversity and Non-African Admixture. PLOS Genetics 8:e1003080.

Pospisilik JA, Schramek D, Schnidar H, Cronin SJF, Nehme NT, Zhang X, Knauf C, Cani PD, Aumayr K, Todoric J, et al. 2010. Drosophila Genome-wide Obesity Screen Reveals Hedgehog as a Determinant of Brown versus White Adipose Cell Fate. Cell 140:148–160.

Quinlan AR. 2014. BEDTools: The Swiss-Army Tool for Genome Feature Analysis. Current Protocols in Bioinformatics 47:11.12.1-11.12.34.

Rai M, Coleman Z, Curley M, Nityanandam A, Platt A, Robles-Murguia M, Jiao J, Finkelstein D, Wang Y- D, Xu B, et al. 2021. Proteasome stress in skeletal muscle mounts a long-range protective response that delays retinal and brain aging. Cell Metabolism 33:1137–1154.e9.

Ramnarine TJS, Grath S, Parsch J. 2022. Natural variation in the transcriptional response of Drosophila melanogaster to oxidative stress. G 3 Genes |Genomes|Genetics 12:jkab366.

Reddy BA, Bajpe PK, Bassett A, Moshkin YM, Kozhevnikova E, Bezstarosti K, Demmers JAA, Travers AA, Verrijzer CP. 2010. Drosophila Transcription Factor Tramtrack69 Binds MEP1 To Recruit the Chromatin Remodeler NuRD. Molecular and Cellular Biology 30:5234–5244.

Rhee DY, Cho D-Y, Zhai B, Slattery M, Ma L, Mintseris J, Wong CY, White KP, Celniker SE, Przytycka TM, et al. 2014. Transcription Factor Networks in Drosophila melanogaster. Cell Reports 8:2031– 2043.

Robinson MD, McCarthy DJ, Smyth GK. 2010. edgeR: a Bioconductor package for differential expression analysis of digital gene expression data. Bioinformatics 26:139–140.

Ross-Innes CS, Stark R, Teschendorff AE, Holmes KA, Ali HR, Dunning MJ, Brown GD, Gojis O, Ellis IO, Green AR, et al. 2012. Differential oestrogen receptor binding is associated with clinical outcome in breast cancer. Nature 481:389–393.

Schuettengruber B, Cavalli G. 2009. Recruitment of Polycomb group complexes and their role in the dynamic regulation of cell fate choice. Development 136:3531–3542.

Schuettengruber B, Ganapathi M, Leblanc B, Portoso M, Jaschek R, Tolhuis B, van Lohuizen M, Tanay A, Cavalli G. 2009. Functional Anatomy of Polycomb and Trithorax Chromatin Landscapes in Drosophila Embryos. PLOS Biology 7:e1000013.

Schuettengruber B, Martinez A-M, Iovino N, Cavalli G. 2011. Trithorax group proteins: switching genes on and keeping them active. Nat Rev Mol Cell Biol 12:799–814.

Schwartz YB, Kahn TG, Stenberg P, Ohno K, Bourgon R, Pirrotta V. 2010. Alternative Epigenetic Chromatin States of Polycomb Target Genes. PLOS Genetics 6:e1000805.

Shingleton AW, Estep CM, Driscoll MV, Dworkin I. 2009. Many ways to be small: different environmental regulators of size generate distinct scaling relationships in Drosophila melanogaster. Proceedings of the Royal Society B: Biological Sciences 276:2625–2633.

Shokri L, Inukai S, Hafner A, Weinand K, Hens K, Vedenko A, Gisselbrecht SS, Dainese R, Bischof J, Furger E, et al. 2019. A Comprehensive Drosophila melanogaster Transcription Factor Interactome. Cell Reports 27:955–970.e7.

Sigrist CJA, Pirrotta V. 1997. Chromatin Insulator Elements Block the Silencing of a Target Gene by the Drosophila Polycomb Response Element (PRE) but Allow trans Interactions Between PREs on Different Chromosomes. Genetics 147:209–221.

Simon JA, Kingston RE. 2013. Occupying Chromatin: Polycomb Mechanisms for Getting to Genomic Targets, Stopping Transcriptional Traffic, and Staying Put. Molecular Cell 49:808–824.

Steffen PA, Ringrose L. 2014. What are memories made of? How Polycomb and Trithorax proteins mediate epigenetic memory. Nature Reviews Molecular Cell Biology 15:340–356.

Stephan W, Li H. 2007. The recent demographic and adaptive history of Drosophila melanogaster. Heredity 98:65–68.

Stojnic R, Diez D. 2022. PWMEnrich: PWM enrichment analysis. Available from: https://bioconductor.org/packages/PWMEnrich/

Svetec N, Werzner A, Wilches R, Pavlidis P, Álvarez-Castro JM, Broman KW, Metzler D, Stephan W. 2011. Identification of X-linked quantitative trait loci affecting cold tolerance in Drosophila melanogaster and fine mapping by selective sweep analysis. Molecular Ecology 20:530–544.

Trotta V, Calboli FC, Ziosi M, Guerra D, Pezzoli MC, David JR, Cavicchi S. 2006. Thermal plasticity in Drosophila melanogaster: A comparison of geographic populations. BMC Evol Biol 6:67.

Ugrankar R, Berglund E, Akdemir F, Tran C, Kim MS, Noh J, Schneider R, Ebert B, Graff JM. 2015. Drosophila glucome screening identifies Ck1alpha as a regulator of mammalian glucose metabolism. Nature Communications 6:7102.

Van Bortle K, Nichols MH, Li L, Ong C-T, Takenaka N, Qin ZS, Corces VG. 2014. Insulator function and topological domain border strength scale with architectural protein occupancy. Genome Biology 15:R82.

Vidal M. 2014. Polycomb Complexes: Chromatin Regulators Required for Cell Diversity and Tissue Homeostasis. In: Bonifer C, Cockerill PN, editors. Transcriptional and Epigenetic Mechanisms Regulating Normal and Aberrant Blood Cell Development. Epigenetics and Human Health. Berlin, Heidelberg: Springer. p. 95–139. Available from: https://doi.org/10.1007/978-3-642-45198-0_5

Voigt S, Erpf AC, Stephan W. 2019. Decreased Temperature Sensitivity of Vestigial Gene Expression in Temperate Populations of Drosophila melanogaster. Genes 10:498.

Voigt S, Kost L. 2021. Differences in temperature-sensitive expression of PcG-regulated genes among natural populations of Drosophila melanogaster. G 3 Genes |Genomes|Genetics 11:jkab237.

Voigt S, Laurent S, Litovchenko M, Stephan W. 2015. Positive Selection at the Polyhomeotic Locus Led to Decreased Thermosensitivity of Gene Expression in Temperate Drosophila melanogaster. Genetics 200:591–599.

Wang L, Brown JL, Cao R, Zhang Y, Kassis JA, Jones RS. 2004. Hierarchical recruitment of polycomb group silencing complexes. Mol Cell 14:637–646.

Wang L, Jahren N, Miller EL, Ketel CS, Mallin DR, Simon JA. 2010. Comparative Analysis of Chromatin Binding by Sex Comb on Midleg (SCM) and Other Polycomb Group Repressors at a Drosophila Hox Gene. Molecular and Cellular Biology 30:2584–2593.

Weir BS, Cockerham CC. 1984. Estimating F-Statistics for the Analysis of Population Structure. Evolution 38:1358–1370.

Wu T, Hu E, Xu S, Chen M, Guo P, Dai Z, Feng T, Zhou L, Tang W, Zhan L, et al. 2021. clusterProfiler 4.0: A universal enrichment tool for interpreting omics data. The Innovation 2:100141.

Zhang Y, Liu T, Meyer CA, Eeckhoute J, Johnson DS, Bernstein BE, Nusbaum C, Myers RM, Brown M, Li W, et al. 2008. Model-based analysis of ChIP-Seq (MACS). Genome Biol 9:R137.

Xie X-J, Hsu F-N, Gao X, Xu W, Ni J-Q, Xing Y, Huang L, Hsiao H-C, Zheng H, Wang C, et al. 2015. CDK8-Cyclin C Mediates Nutritional Regulation of Developmental Transitions through the Ecdysone Receptor in Drosophila. PLOS Biology 13:e1002207.

Yao L, Wang S, Westholm JO, Dai Q, Matsuda R, Hosono C, Bray S, Lai EC, Samakovlis C. 2017. Genome-wide identification of Grainy head targets in Drosophila reveals regulatory interactions with the POU domain transcription factor Vvl. Development 144:3145–3155.

Zhao L, Wit J, Svetec N, Begun DJ. 2015. Parallel Gene Expression Differences between Low and High Latitude Populations of Drosophila melanogaster and D. simulans. PLOS Genetics 11:e1005184.

Zink D, Paro R. 1995. Drosophila Polycomb-group regulated chromatin inhibits the accessibility of a trans-activator to its target DNA. The EMBO Journal 14:5660–5671.

