## Supplementary Table S38 for "Genome-wide temperature-sensitivity of PcG regulation and reduction thereof in temperate *Drosophila melanogaster*"

Table S38. Transcription factors with motif differentiation between populations, overrepresentation at PcG target genes with population x temperature interactions in their expression, and known protein-protein interactions with PcG recruiter, PcG, TrxG and insulator proteins.

| Transcription factor | Interactions | References |
|---|---|---|
| bap | insulator (BEAF-32) | Shokri et al. (2019) |
| bcd | TrxG (nej) | Fu et al. (2004) |
| br | PcG (Pc) | Lv et al. (2016) |
| da | TrxG (Act42A, Act5C) | Hsiao et al. (2014) |
| dl | PcG recruiter (Dsp1, Spps); TrxG (d4, nej) | Lehming et al. (1994); Decoville et al. (2000); Holmqvist et al. (2012); Shokri et al. (2019) |
| Dll | PcG recruiter (Dsp1) | Hu et al. (2018) |
| E(spl)m3-HLH | TrxG (Snr1) | Guruharsha et al. (2011) |
| EcR | insulator (Chro, Cp190, CTCF); PcG recruiter (cg, Psq); TrxG (brm, E(bx), Lpt, MBD-like, Mi-2, mor, nej, osa, trr, Utx) | Badenhorst et al. (2005); Chauhan et al. (2012); Denton et al. (2013); Hitrik et al. (2016); Kreher et al. (2017); Pascual-Garcia et al. (2017); Gutierrez-Perez et al. (2019); Shokri et al. (2019); Mazina et al. (2020) |
| Eip78C | insulator (Clamp); PcG (pho; ph-p); PcG recruiter (pho, grh); TrxG (d4, MDB-like) | Shokri et al. (2019) |
| en | TrxG (nej) | Luo et al. (2017) |
| Ets96B | insulator (su(Hw)) | Shokri et al. (2019) |
| fru | TrxG (HDAC1) | Ito et al. (2012) |
| GATAe | insulator (M1BP, ZIPIC) | Shokri et al. (2019) |
| Gsc | insulator (BEAF-32, Ibf2) | Hu et al. (2018) |
| h | PcG (phol); PcG recruiter (phol) | Shokri et al. (2019) |
| Hr78 | insulator (Clamp, CTCF, M1BP, su(HW), Trl, ZIPIC); PcG (esc, pho, ph-p); PcG recruiter (grh, pho, Trl); TrxG (ash2, d4, HDAC1) | Shokri et al. (2019) |
| Kr | TrxG (d4) | Shokri et al. (2019) |
| lola | insulator (Trl); PcG recruiter (Trl) | Lomaev et al. (2017) |
| Mad | PcG (Scm); TrxG (nej) | Lilja et al. (2007); Shokri et al. (2019) |
| retn | PcG (pho); PcG recruiter (pho, Spps) | Shokri et al. (2019) |
| sens-2 | PcG (phol); PcG recruiter (phol) | Shokri et al. (2019) |
| sv | PcG (phol); PcG recruiter (phol) | Shokri et al. (2019) |
| ttk | insulator (Trl); PcG recruiter (Trl); TrxG (CDK2AP1, MBD-like, MEP-1, Mi-2, simj) | Pagans et al. (2002); Pagans et al. (2004); Reddy et al. (2010); Bonchuk et al. (2011); Rhee et al. (2014); Khalisova et al. (2021) |
| usp | TrxG (Lpt, Utx) | Chauhan et al. (2012); Denton et al. (2013); Xie et al. (2015) |
| vis | PcG recruiter (grh) | Shokri et al. (2019) |
| vvl | PcG recruiter (grh) | Yao et al. (2017) |
| wor | TrxG (MBD-like) | Shokri et al. (2019) |
